# Machine learning and data-driven inverse modeling of metabolomics unveil key process of active aging

**DOI:** 10.1101/2024.08.27.609825

**Authors:** Jiahang Li, Martin Brenner, Iro Pierides, Barbara Wessner, Bernhard Franzke, Eva-Maria Strasser, Steffen Waldherr, Karl-Heinz Wagner, Wolfram Weckwerth

**Affiliations:** Molecular Systems Biology Lab (MOSYS), Department of Functional and Evolutionary Ecology), University of Vienna, Djerassiplatz 1, 1030 Vienna, Austria; Vienna Molecular Metabolomics Center (VIME), University of Vienna, Djerassiplatz 1, 1030 Vienna, Austria; Department of Nutritional Sciences, University of Vienna, Vienna, Austria; Research Platform Active Ageing, University of Vienna, Josef-Holaubek-Platz, 1090 Vienna

**Keywords:** Active aging, metabolic network inference, automatic machine learning, data-driven modeling

## Abstract

Physical inactivity and a weak fitness status have become a global health concern. Metabolomics, as an integrative systematic approach, might link to individual’s fitness at the molecular level. In this study, we performed blood samples metabolomics analysis of a cohort of elderly people with different treatments. By defining two groups of fitness and corresponding metabolites profiles, we tested several machine learning classification approaches to identify key metabolite biomarkers, which showed robustly aspartate as a dominant negative marker of fitness. Following, the metabolomics data of the two groups were analyzed by a novel approach for metabolic network interaction termed COVRECON. Where we identified the enzyme AST as the most important metabolic regulation between the fit and the less fit groups. Routine blood tests in these two cohorts validated significant differences in AST and ALT. In summary, we combine machine learning classification and COVRECON to identify metabolomics biomarkers and causal processes for fitness of elderly people.

## Introduction

Physical inactivity is a worldwide health problem, and is ranked as the fourth leading behavioral risk factor for global mortality (Kohl, et al., 2012). The imperative to maintain body activity, physically and metabolically is on the rise. The concept of active aging, inspired by Robert Havighurst’s activity theory (Havighurst, 1963), suggests that maintaining an active lifestyle is crucial for the well-being of older individuals. The thought of active aging emerged and began to develop in the 1990s, placing strong emphasis on the link between activity and health (WHO, 2002).This focus became particularly pertinent due to the worldwide aging population, leading to concerns about inactivity permeating various social domains (Boudiny and Mortelmans, 2011). Within the transition into the 2020s, there has been an escalating emphasis on harnessing technology to foster healthy aging (Malkowski, et al., 2023; Offerman, et al., 2023; Wongsala, et al., 2021). Beyond longevity, active aging encompasses regular physical activity, better management of chronic diseases, and improved quality of life (Fernández-Ballesteros, et al., 2013).

Conventionally, several studies have examined various physical aspects of active aging, such as sleep, sedentary time, muscle strengthening activities, and movement behaviors (Caprara, et al., 2013; Taylor, 2022). As the new technology to diagnose diseases, metabolomics involves the comprehensive analysis of small-molecule metabolites (<10 kDa) present in a biological sample, including metabolic intermediates, hormones, signaling molecules, and secondary metabolites (Patti, et al., 2012; Weckwerth, 2010). Functioning as the culmination of all biological processes in the body, metabolites play a pivotal role in energy generation, signal transmission, and carrying essential information about the body’s status and ongoing functions. Consequently, important metabolites possess the potential to serve as aging biomarkers or as integral components of the metabolic signature. This signature mirrors the active state of the organism as it traverses the aging process (Balashova, et al., 2022). The development of metabolomics empowers us to scrutinize health issues at the molecular level (Gonzalez-Covarrubias, et al., 2022). Notably, amid the COVID-19 pandemic, metabolomics has demonstrated its potency as a diagnostic, prognostic, and drug intervention tool (Bruzzone, et al., 2023). As expected, COVID-19 has been extensively investigated using metabolomics methodologies, contributing to biomarker studies (Sindelar, et al., 2021; Su, et al., 2020) and evaluations of drug impacts (Ghini, et al., 2022; Meoni, et al., 2021). Beyond specific disease diagnosis, metabolomics can also illuminate our comprehension of bodily activities (active aging). Recent endeavors have delved into metabolic profiling within aging studies (Balashova, et al., 2022; Panyard, et al., 2022), providing us with overarching insights into metabolic changes during the aging process. Here we specifically focus on the metabolomics profiling of older adults and aim to detect key biomarkers and important metabolic interactions related to active aging.

The emerging large scale datasets from OMICS (metabolomics, proteomics, transcriptomic and genomics) measurements empower us to scrutinize any question in biology from a systemic perspective (Weckwerth, 2011; Weckwerth, 2019). In the field of systems biology, a central goal is to identify biomarkers and infer biochemical regulations from large-scale metabolomics data (Weckwerth, 2011). Statistical methods, especially when combined with machine learning techniques, have shown power on constructing accurate classifiers capable of distinguishing between diverse sample groups and revealing underlying biomarkers (Alakwaa, et al., 2018; Liebal, et al., 2020; Pomyen, et al., 2020; Sidak, et al., 2022). However, statistical methods offer limited insights into how information is transferred within a biochemical network, the critical regulatory steps involved, and how regulatory mechanisms change under different conditions (Weckwerth, 2003; Weckwerth, 2010; Weckwerth, 2011; Weckwerth, 2019). Several studies have emphasized the necessity of analyzing the dynamic behavior of metabolism to understand the evolution and maintenance of stable metabolic homeostasis under varying environmental conditions (Li, et al., 2023; Nägele, et al., 2014; Wienkoop, et al., 2008; Wilson, et al., 2020).

Mathematically, kinetic models can provide systemic insights into metabolic networks, but constructing these models and estimating parameters poses challenges, particularly for large-scale models (King, et al., 2016). In recent years, several studies have focused on the steady-state Jacobian investigation of metabolomics data (Akbari, et al., 2024; Haiman, et al., 2021; Jamshidi and Palsson, 2010; Nägele, 2022; Steuer, et al., 2006) integrating with fluxomics or time-series measurements. In addition, with only large sampled metabolomics measurements, recent studies have developed inverse differential Jacobian algorithms, which provide a convenient way to infer differences in the dynamics of metabolic networks between different conditions (Kügler and Yang, 2014; Li, et al., 2023; Nägele, et al., 2014; Sun, et al., 2015; Sun and Weckwerth, 2012; Weckwerth, 2019; Weiszmann, et al., 2023; Wilson, et al., 2020). Among them, the most recent study has a novel method and workflow termed COVRECON for analyzing key biochemical regulations through the solution of a differential Jacobian problem (Weckwerth, 2019; Li, et al., 2023). The COVRECON approach integrates the covariance matrix of metabolomics data with automatic metabolic network modeling based on genome-scale metabolic reconstructions and biochemical reaction databases.

Figure 1 illustrates the workflow of this study, which consists of three main steps. In step 1, we aim to cluster the original samples into different groups based on physical and functional measurements, where each group represents different body activity conditions. Step 2 involves building machine learning-based classifiers to identify these different groups using metabolomics data, thus enabling the identification of key metabolites as biomarkers. Finally, in step 3, we employ the inverse Jacobian analysis and the COVRECON workflow to uncover the most important biochemical regulations associated with the identified body activity conditions. By conducting this approach, we intend to contribute to the understanding of active aging from a metabolomics perspective and shed light on the key biochemical regulations underlying different fitness conditions.

**Figure 1:**
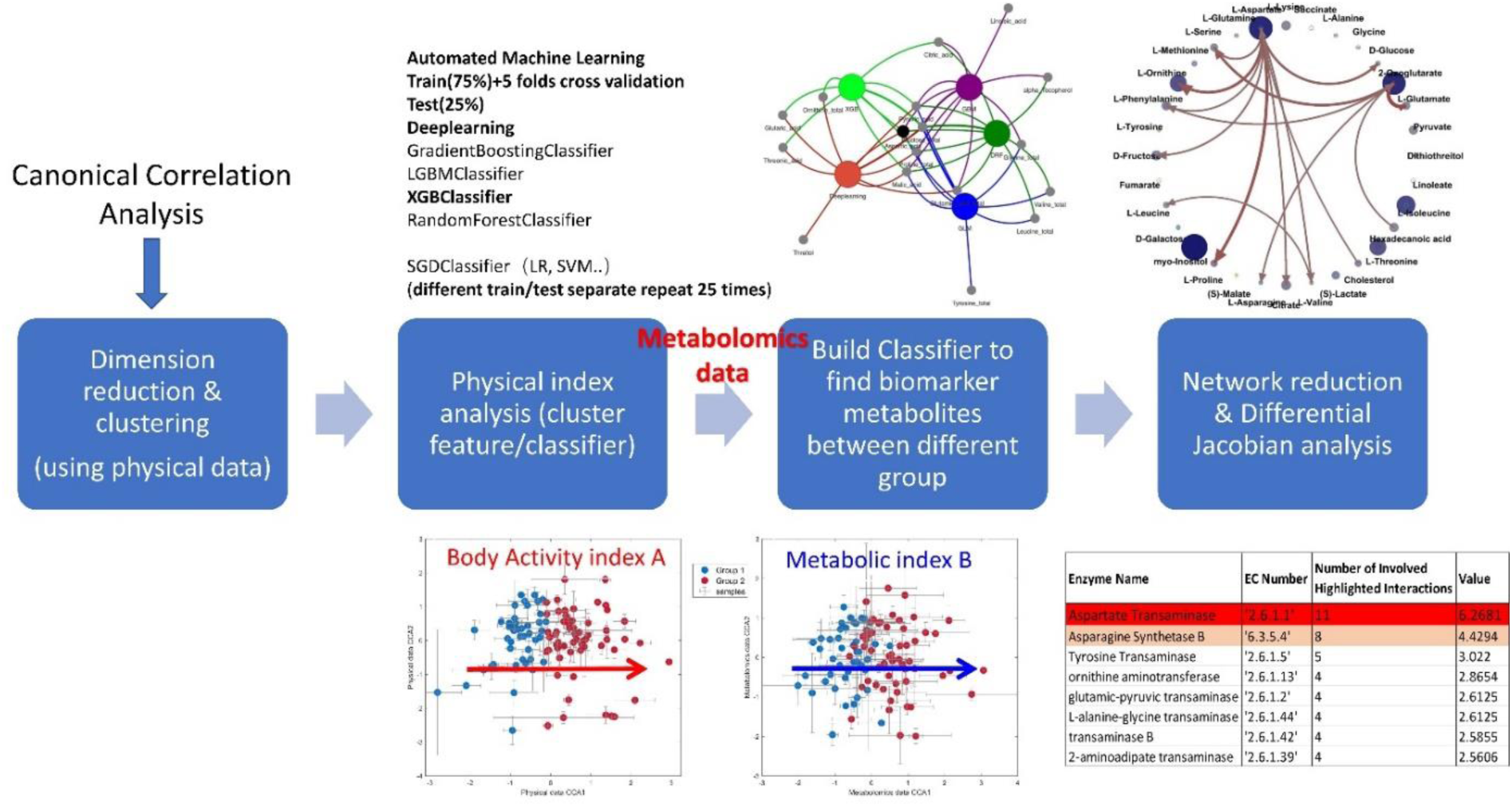
Work scheme of the proposed approach. This figure illustrates the key steps of our proposed methodology. In the first stage, we employ the Canonical Correlation Algorithm (CCA) to derive a body activity index that is highly linked to metabolomics data. Subsequently, we perform sample clustering, wherein samples are grouped into 2, 4, and 6 clusters based on this index. To address the non-linear effects of metabolites, we develop a set of automated machine-learning classifiers specifically tailored to metabolomics data, allowing us to identify crucial variables. Finally, we employ the COVRECON workflow (Li, 2023) to construct a topological metabolic interaction model for measured metabolites and to solve the differential Jacobian problem of the system.

## Materials and Methods

### Experimental design

This study was performed in 5 retirement homes in Vienna managed by Curatorship of Viennese Retirement Homes. The aim of this study was to assess the impact of strength training, strength training and protein-vitamin supplement or a cognitive training on very old, institutionalized adults. This study was conducted in a randomized, controlled, observer-blind design. The subjects were randomly assigned to three groups: resistance training (T), resistance training and supplements (E) and cognitive training, acting as a control group (K). The details are presented in Supplementary material S1. Blood samples were collected at the baseline (T1), after three months (T2) and after six months (T3).

One hundred seventeen subjects were recruited from five senior residencies (Supplementary Figure S1). The exclusion criteria consisted of physical fitness (Short Physical Performance Battery > 4) and mental performance (Mini Mental State Examination ≥ 23). Moreover, they were free of severe diseases such as diabetic retinopathy, CVDs and regular use of cortisone-containing drugs. Before starting the intervention the health and nutritional status was assessed by specialists in internal medicine and gerontology (Oesen, et al., 2015). All subjects signed informed consent before inclusion in accordance with the Declaration of Helsinki. The study was approved by the ethics committee of the City of Vienna (EK-11-151-0811) and registered at ClinicalTrials.gov, NCT01775111 (Oesen, et al., 2015).

### Subject characteristics

The sex distribution (87.6% women; 12.4% men) among participants was representative for the population living in nursing homes. The mean age of the study population was 82.9± 6.0 years for women and 84.9 ± 6.7 years for men. The participants had a BMI of 29.27 kg/m^2^ ±5.00 kg/m^2^ (Oesen, et al., 2015).

### Sample Preparation

#### Blood Plasma Metabolite Extraction

Several studies addressed the choice of blood sample, revealing that Heparin plasma produces a smaller side effect in the chromatogram spectrum (Dunn, et al., 2011; Teahan, et al., 2006). Concordant with these findings, Heparin was used as an anticoagulant, while blood plasma was separated from fresh blood samples and kept in −80° C for further clinical analysis. Metabolite profiles of obtained human plasma samples was measured using a gas chromatograph coupled to mass spectrometer (Weckwerth, et al., 2004). The samples were thawed on ice for 45 min and were vigorously vortexed for 10 s. The extraction consisted of two steps. First, 100µl plasma were transferred into 1.5 ml Eppendorf tubes, followed by the addition of 600 µl ice cooled MeOH, immediately vortexed for 10 s and left one ice for 15 min for incubation. In order to remove proteins, the samples were centrifuged at 14000 g for 4 min at 4° C. The supernatant was transferred into new tubes and dried down in a SpeedVac. Afterwards the dried pellets were stored at −20°C.

The second step consisted of extraction with CHCl3. 300 µl of CHCl3 were added to pellets. The further procedure was a repetition of the first step. The supernatant was transferred into new Eppendorf tubes and dried down in SpeedVac. Metabolite extractions were performed in batches of 30 samples, of randomly selected subjects.

#### Quality control-Mix

Quality control (QC) consisted of specific metabolites, including organic acids, amino acids, mono- and disaccharides and substrates of the TCA cycle. The table of metabolites for QC-Mix is attached to supplemental material S2. A calibration curve was prepared with concentrations of 2 µl, 5 µl, 10 µl, 20 µl, 40 µl, 80 µl and 100 µl.

Internal Standard (10 µl Pinitol and 10 µl Sorbitol) was added to each sample and to each QC just the day before GC-MS analysis. Afterwards, the samples were dried in a SpeedVac.

#### Derivation

First, addition of 20 µl of 40 mg mL^-1^ of methoxyamine hydrochloride (MeOX) dissolved in pyridine were added to each sample in order to dissolve MeOX in pyridine appropriately, the solution was vigorously vortexed several times and tube was put into hot water. After that, samples were vortexed until pellets were completely dissolved, followed by agitation at 30 °C for 90 min at 750 rpm with a thermoshaker.

N-Methyl-N-(trimethylsilyl) trifluoroacetamide (MSTFA) flasks of 1 ml content were spiked with 30 µl retention index marker solution of alkanes from C10-C40 in hexane. After addition of 80 µl of prepared MSTFA, samples were incubated at 37° C for 30 min at 750 rpm, followed by centrifugation at 14000 g for 2 min at room temperature (24° C). Immediately after this step, 70 µl of the supernatant were transferred to GC-vials with micro inserts and closed with crimp caps.

#### GC-MS Analysis

Finally, samples were analysed using GC-MS (LECO Pegasus® 4D GCxGC-TOF-MS, Mönchengladbach, Germany) according to Weckwerth, Wenzel, and Fiehn 2004. Immediately after derivation, 1 µl of sample were injected utilizing a split ratio of 1:5. The split/splitless injector was kept at a constant temperature of 230°C equipped with a single-tapered liner with deactivated wool. The GC-MS consisted of an Agilent 6890 (Agilent Technologies, Glostrup, Denmark) using helium as carrier gas at a flow rate of 1 mL min–1. Gas separation was performed on the HP-5MS column (30 m 3 0.25 mm 3 0.25 mm, Agilent Technologies).

The initial temperature of the GC oven was set to 70°C isothermal for 1 min, followed by a heating ramp of 9°C min^−1^ to reach 330°C and hold for 7 min.

Transfer line temperature was 250°C, and ion source temperature was set to 200°C. The MS detector was switched off during the first 260 sec. Mass spectra were acquired with an acquisition rate of 20 spectra s^−1^ and were recorded in the range of 40 to 600 m/z, utilizing a detector voltage of 1,550 V and electron impact ionization of 70 eV. The metabolite assessment required an exchange of the liner every 70 injections, thus every 2 batches in a row.

The whole data acquisition was performed within 14 batches. Each batch was measured in the same chronological order. First, alkanes were injected, followed by QC calibration curve. Blank sample that contained only dried extraction reagents and derivation solvents were injected each 5 or 7 samples. Each batch consisted of a single plasma sample from 20-30 subjects and was analyzed within 24 – 32 hours. One pooled sample was measured for each batch, in order to assess instrument stability. At the end of every batch, the same QC was measured again to monitor instrumental performance over time. To minimize systematic bias induced by preparation order, samples were randomly distributed into the 14 batches. However, each batch consists of a representative cross section of total samples and was comparable to the total experimental population.

#### Metabolite identification

Mass spectra data were obtained from GC-MS, next step is to transform this to biologically relevant information.

After the GC-MS analysis the raw data consisted of ion peaks and were preprocessed using LECO Chroma-TOF. The ion fragmentation spectra were matched to fragmentation spectra in NIST library and scored with a match probability, taking into account only metabolites with at least 700 similarity score. Afterwards, analytes were identified by comparison of ion fragments to a reference library of chemical standards. In detail, the metabolites were confirmed based on ion features e.g. 1) retention index and retention time, 2) m/z, 3) in-source fragmentation, particular for each metabolite. With the latter the identification of the analytes became definitive.

Alkanes measured at the beginning of each batch provided retention indices that were assigned to all ion peaks.

The original data is presented in Supplementary material S2.

#### Data Processing

The data processing steps involved several procedures. Initially, missing values in the metabolomics measurements were imputed using the K-Nearest Neighbors (KNN) method. Following, normalization was performed to reduce heteroscedasticity and adjust for the offset between high and low-intensity features as in Equation (0), where the log transformation of each metabolite by centering it around its mean (X̅) and scaling it by its standard deviation (s):

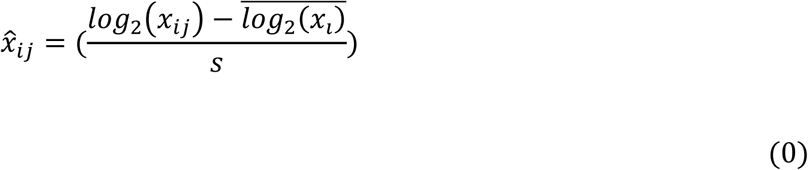

#### Data Clustering

To identify biomarkers and perform the inverse Jacobian analysis, the samples were firstly clustered into distinct groups. The clustering process comprised the following steps. Firstly, based on the information provided in Supplementary Material S2, it was observed that physical measurements could be categorized into two types: “body-shape” data (e.g., gender and height) and “body-functional” data (e.g., walking distance and left standing time). In order to generate a body activity index that reflects body functionality while minimizing the influence of body-shape differences, Canonical Correlation Analysis (CCA) (Hardoon, et al., 2004) was applied. The loadings of this body activity index are presented in Figure 2A, where it can be observed that walking distance exhibits the strongest effects. The metabolomics-related body activity index generated through CCA was then used to cluster the samples using the k-means method, grouping them based on this body activity index.

**Figure 2.**
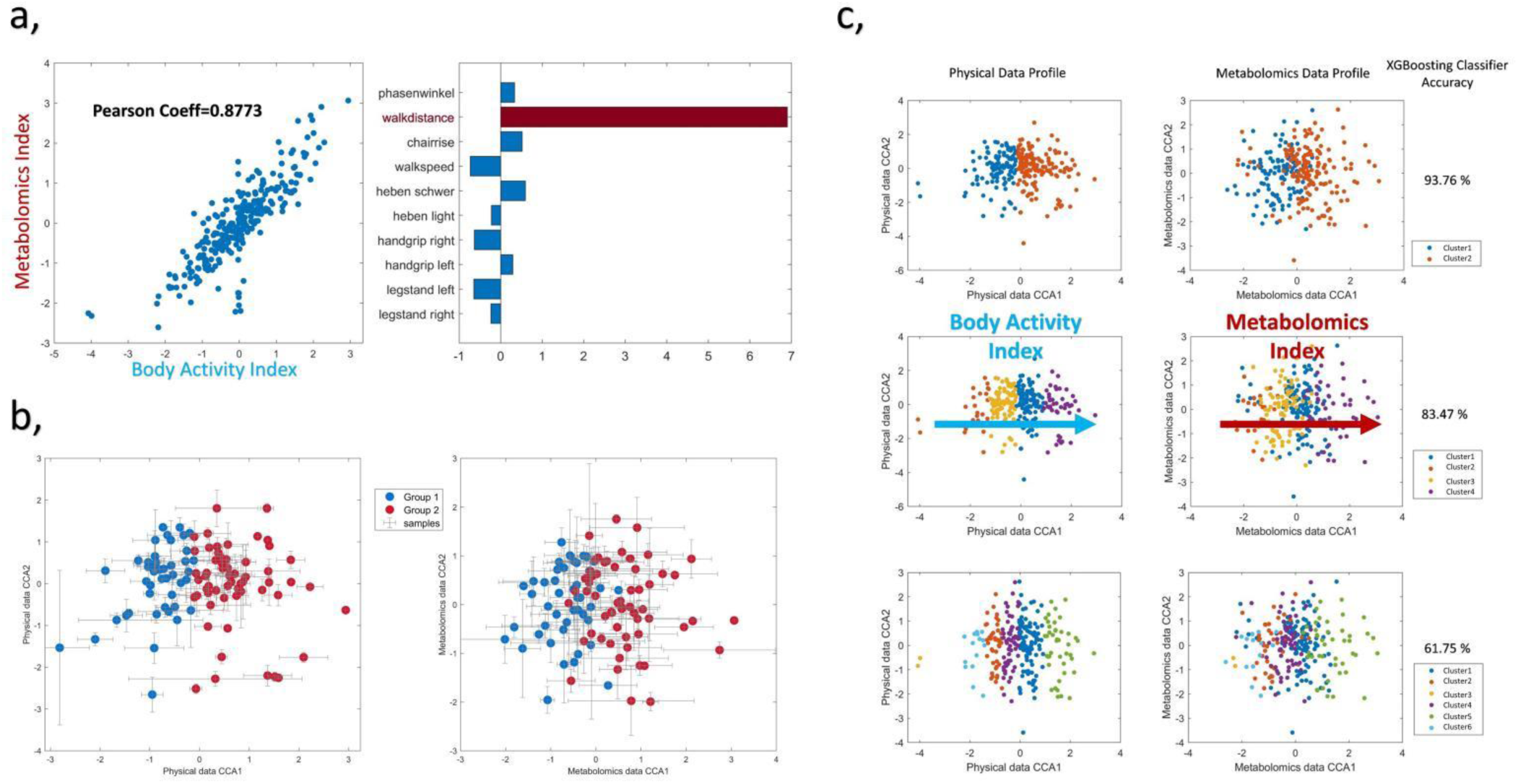
Body Activity Index and Metabolomics Index from canonical corresponding analysis (CCA) and cluster analysis. (a), The scatter plot of the generated Body Activity Index and Metabolomics Index and loadings of Body Activity Index. (b), The scatter plots of physical (left) and metabolomics (right) profiles, where all old adults are clustered into two groups based on Body Activity Index. (c), The scatter plots of physical (left) and metabolomics (right) profiles with all samples clustered into 2, 4 and 6 groups.

#### Machine Learning based Classifiers

While the CCA-based clustering approach analyzes the relationship between the body activity index and the metabolic index as a linear method, it may not fully capture the dynamic nature of the metabolic mechanism, which inherently exhibits predominantly non-linear behavior. To capture this non-linear influence and achieve higher accuracy with the identification of important variables, several machine learning based classifiers were employed within an automated machine learning framework, implemented using the H2o package in Python. The methods utilized are as follows:

1, Generalized Linear Models (GLM): GLM implements regularized linear models with stochastic gradient descent (SGD) learning. The model is updated iteratively using a decreasing strength schedule, estimating the loss gradient for each sample at a time. This method offers a baseline for the linear effects.
2, Random Forest Classifier: A random forest is an ensemble meta-estimator that fits multiple decision tree classifiers on different sub-samples of the dataset, utilizing averaging to improve predictive accuracy and mitigate overfitting.
3-4, Boosting Methods: Boosting is an ensemble meta-algorithm that reduces bias and variance in supervised learning. It integrates a family of machine learning algorithms that convert weak learners to strong ones (Zhou, 2012). The main variation between many boosting algorithms is their method of weighting training data points and hypotheses. We employed two common boosting methods, LGBMClassifier and XGBClassifier. LGBMClassifier (GBM) is a distributed gradient-boosting framework based on decision tree algorithms, originally developed by Microsoft (Ke, et al., 2017), while XGBClassifier (eXtreme Gradient Boosting) is an open-source library for regularizing gradient boosting (Chen and Guestrin, 2016).
5, Autoencoder + deep learning: Deep learning, also known as deep neural networks, is a powerful machine learning method extensively used in pattern recognition, image processing, and bioinformatics (LeCun, et al., 2015). Prior to training the model, we employed an autoencoder to pre-train it, using the entire unlabeled data, improving model performance, preventing random weight initialization.

In our approach, each of these machine learning methods was integrated into an automated framework that encompasses hyper-parameter optimization. Hyper-parameter optimization entails the selection of ideal parameter values that govern the learning process, aiming to enhance model performance (Feurer and Hutter, 2019). The supplementary Figure S5 provides an overview of the scope of hyper-parameters associated with each machine learning method. For evaluation, we randomly generate 25 training-test separations where the training-test ratio is 75/25 %.

#### Feature Importance

Feature importance was estimated using a model-based approach, considering a feature to be important if it significantly contributed to the model’s performance. Here, the ‘varimp’ function within the H2o.py package was utilized to rank the important metabolites of each classifier. The importance value is averaged over the 25 training-test separations, and we choose the top 10 metabolites for each machine-learning method.

#### Predictive metabolic modelling using an inverse Jacobian approach

Statistical and machine learning methods have limitations when it comes to understanding the dynamics of a biochemical network, identifying critical regulatory steps, and capturing changes in regulatory mechanisms under different conditions (Sidak, et al., 2022). In recent years, the inverse differential Jacobian algorithms have been developed as a convenient approach to infer the dynamic regulation of metabolic networks from metabolomics data (Kügler and Yang, 2014; Li, et al., 2023; Nägele, et al., 2014; Steuer, et al., 2003; Sun, et al., 2015; Sun and Weckwerth, 2012; Weckwerth, 2019; Wilson, et al., 2020).

In previous studies, we introduced the COVRECON workflow and Matlab toolbox as the standard inverse Jacobian workflow (Weckwerth, 2019; Li, et al., 2023). This method combines the covariance matrix of metabolomics data with automatic metabolic network modeling based on genome-scale metabolic reconstructions and biochemical reaction databases.

Consider a metabolic network that consists of n metabolites denoted by {*X*_*i*_}_*i*=1…*n*_. The system dynamics can be modeled with the set of ordinary differential equations (ODEs):

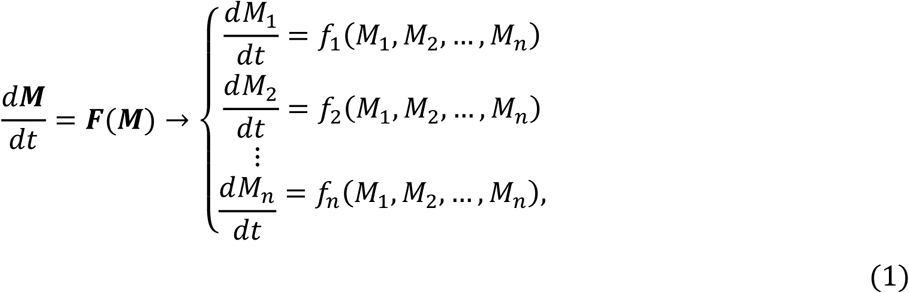

where *M* = {*M*_*i*_} = {|*X*_*i*_|} are the concentrations of the n metabolites, and *F* = *f*_*i*_(*M*_*i*_) are composed of the reaction rates for these metabolites (e.g., Michaelis-Menten kinetics, or mass action).

The steady-state Jacobian matrix *J* of the model is defined as a *R*^*n*×*n*^ matrix in which *J*_*ij*_ is the first-order derivative of the rate *f*_*i*_ for the concentration of substances *M*_*j*_ at steady state, noted as 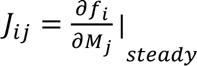:

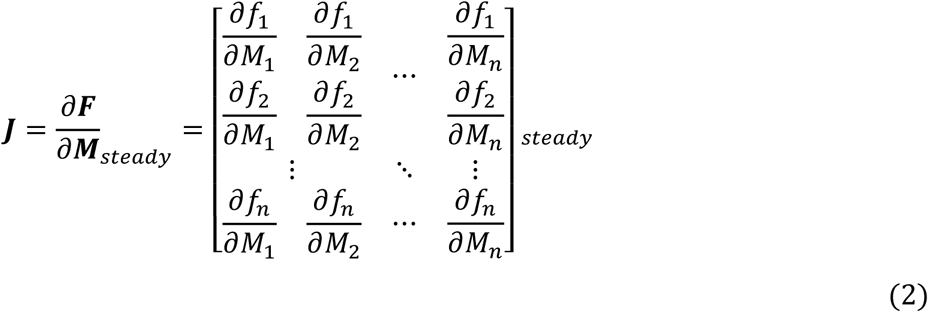

The steady-state Jacobian matrix represents the first-order derivatives of the rate equations with respect to the concentrations of the metabolites at steady state. It contains valuable information about the system’s dynamics, including regulatory interactions among the metabolites. As derived in a previous study by Steuer et al. (Steuer et al., 2003), the following equation (3) was established between the covariance matrix of the metabolic data, and the steady-state Jacobian matrix:

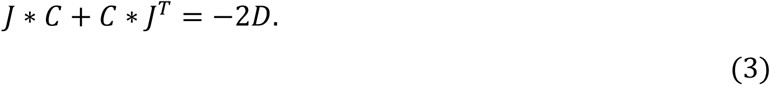

Here, *C* ∈ *R*^*n*×*n*^ represents the covariance matrix of the compounds’ concentrations *M*_*j*_ near its steady-state value 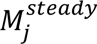, while the fluctuation matrix ***D*** represents the covariance of noise sources acting on the system.

The differences between two conditions can be quantified by the differential Jacobian *D**J***, which is calculated from the Jacobians of the two groups:

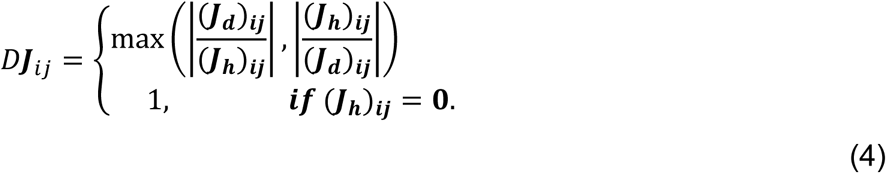

The differential Jacobian *D**J*** encompasses crucial insights into the dynamic regulatory mechanisms between two conditions. An inverse problem is to analyze the differential Jacobian *D**J*** from the measured metabolomics. This task involves two key aspects: establishing the structural information of the Jacobian matrix and resolving the optimization problem associated with the differential Jacobian.

In a recent study, we introduced the COVRECON approach and related matlab toolbox (Li, et al., 2023). This innovative approach combines the automatic assembly of a metabolic interaction network and the inverse differential Jacobian calculation through a regression-loss-based algorithm. This approach automatically constructs a metabolic interaction network which contains the Jacobian structure information and then calculates a regression loss matrix R* to estimate the differential Jacobian matrix. The result R* is presented in a matlab format figure where the interaction pathway details can be interactively checked. The details of the algorithm can be found in the Supplementary material S1 and the original publication (Li, et al., 2023).

By employing the COVRECON approach, we aim to uncover the key components and regulatory interactions within the differential Jacobian, thereby gaining insights into the dynamics of the metabolic network.

#### Integrate Classifier Biomarkers and Group Differential Jacobian Analysis

Since we have clustered the samples into two groups in the data clustering part, we are now able to do the inverse Jacobian analysis for the two groups. As discussed in Supplementary material S1, similar to the general approach of most kinetic models, we consider the dynamics within each group is simulated in a group model, thus the steady state dynamics can be represented as a group Jacobian. Consequently, the inverse Jacobian algorithm can offer valuable information of the regulated dynamics between the two groups.

The results from the inverse Jacobian analysis are closely linked to the structural information of the Jacobian obtained from the automatically generated super-pathway metabolic interaction networks. It is essential to highlight that we combine the significance of classifier variables in the context of inverse Jacobian analysis. Simply put, we retain the pivotal biomarkers and introduce a controlled mix of randomly chosen additional metabolites. The augmented networks, encompassing 10-20 metabolites, are subsequently subjected to the COVRECON workflow. Notably, in COVRECON results, large values serve as indicators of the dynamics difference between the two distinct groups. We are able to identify the important reactions or enzymes involved in the active aging context by checking the detailed information behind these large values (Li, et al., 2023).

## Results

This study was performed in 5 retirement homes in Vienna managed by Curatorship of Viennese Retirement Homes, with the main aim to assess the impact of resistance training and protein-vitamin supplementation or a cognitive training on physical performance (Oesen et al. 2015). In this secondary analysis, we focus on the plasma metabolomics changes and the identification of potential biomarkers and biochemical processes for fitness. The cohort of older adults with an average age above life expectancy consisted of 117 participants at baseline and altogether we measured 263 plasma metabolomic samples.

The subjects were randomly assigned to three groups (see supplementary figure S1): resistance training (T), resistance training and supplements (E) and cognitive training, acting as a control group (K). Blood samples were collected at the baseline (T1), after three months (T2) and after six months (T3).

To establish the relationship between body activity, fitness and metabolomics profiles, we initially investigated the physical data measurements, which consisted of two types: body functionality and body shape. Moreover, the body strength measurements can be further divided into resistance exercise and endurance exercise types. Table 1 shows the group differences of the physical measurements across the three groups. As expected, compared to the control group (K), the resistance training groups (T and E) exhibit better resistance measurements. Nevertheless, there was no influence on endurance measurements (e.g. walking distance). Notably, endurance exercise has been reported more related to body aging conditions than resistance measurements (Cao Dinh, et al., 2019; Werner, et al., 2019). This is also consistent with the experimental design, where the old adults were randomly assigned to the three groups regardless of their fitness.

**Table 1.** , physical measurement difference between three treatment groups.

| physical measurements |  | legstand<br>right | legstand<br>left | handgrip<br>left | handgrip<br>right | heben<br>light | heben<br>schwer | walkspeed | chairrise | walkdistance | phasenwinkel |
| --- | --- | --- | --- | --- | --- | --- | --- | --- | --- | --- | --- |
| group difference<br>t-test p-value | T K difference | 0.8237 | 0.3577 | **0.0003 | **0.0009 | **0.0066 | *0.0124 | 0.6343 | *0.0211 | 0.8551 | **0.0081 |
|  | E K difference | 0.4982 | 0.6228 | **0.0062 | *0.0363 | *0.0146 | *0.0254 | 0.2507 | *0.0109 | 0.8994 | **0.0012 |
|  | E T difference | 0.3638 | 0.1506 | 0.3493 | 0.3993 | 0.9698 | 0.9551 | 0.1024 | 0.3664 | 0.9872 | 0.3414 |

### Canonical Correlation (CCA) based clustering to assess physical fitness in a cohort of older institutionalized adults

Since our question is the relationship between metabolomics and body activity, we employed Canonical Correlation Analysis (CCA) to generate a body activity index based on the body functionality dataset. Subsequently, we clustered the old adults and samples into two, four, and six groups based on this body activity index.

As demonstrated in Figure 2A, the generated body activity index has a high correlation to the metabolomics index (Pearson Coeff = 0.8471), where the CCA loadings of the body activity index is listed aside. Among all physical indexes, walking distance showed the most dominant effect within the body activity index. This observation is biologically reasonable since walking distance directly reflects an individual’s endurance condition, which is directly related to the aging process (Cao Dinh, et al., 2019; Werner, et al., 2019).

Considering potential non-linear relationship between the generated body activity index and the metabolomics index, we constructed an automated machine-learning classifier using the XGBoosting algorithm as described in the method part. The classifier was trained with 100 maximum models, over 25 random training-test separation, the averaged AUCs calculated on the 25% hold-out test sets were determined to be 93.76%, 83.47%, and 61.75% for the two, four, and six-group clusters, respectively (Figure 2C). This indicates that the CCA generated body activity index and metabolomics index exhibit a strong correlation. Meanwhile, we group all the old adults into two groups for the inverse Jacobian analysis using the mean body activity index as shown in Figure 2B.

For comparison, we also performed CCA analysis between the metabolomics data and body shape features, such as gender, height, and age. The biplots of the CCA from the metabolomics and body shape analysis are presented in Supplementary Figure S2. The highest Spearman’s correlation coefficient obtained was only 0.4963 for the age index. Additionally, we conducted a further CCA analysis considering the metabolomics data along with both body functionality and body shape data. However, the Pearson correlation coefficient increased only marginally from 0.8773 to 0.8903. This indicates that the metabolomics data are primarily influenced by the body strength/ functionality aspects. Consequently, this validates both the body activity index and the metabolomics index that we developed.

The results of the CCA-based cluster analysis highlight the strong relationship between the derived body activity index and the metabolomics index. The dominance of walking distance as a key factor indicates its significance as a reflection of an individual’s health condition and metabolic activity. In the following analysis, we will focus on the two old adult groups clustered based on the body activity index, labelled as active group and less-active group.

### Machine Learning based classifiers and variables importance reveals strong association of metabolites and fitness

In this section, we built several machine learning based classifiers to predict the active/less-active groups from the metabolomics dataset. This approach can provide us valuable insights on the nonlinear influence between the metabolomics index and body activity index. As mentioned in the methods part, we evaluated the predicting performance of five machine learning algorithms: XGBoosting, DRF, GLM, GBM, and DeepLearning algorithms. We applied the automatic machine learning framework and selected the best model for each algorithm based on cross-validation AUC values. The chosen models were then assessed on the 25% hold-out test dataset. As shown in Figure 3, the classifiers performances were compared among the five Machine Learning algorithms.

**Figure 3.**
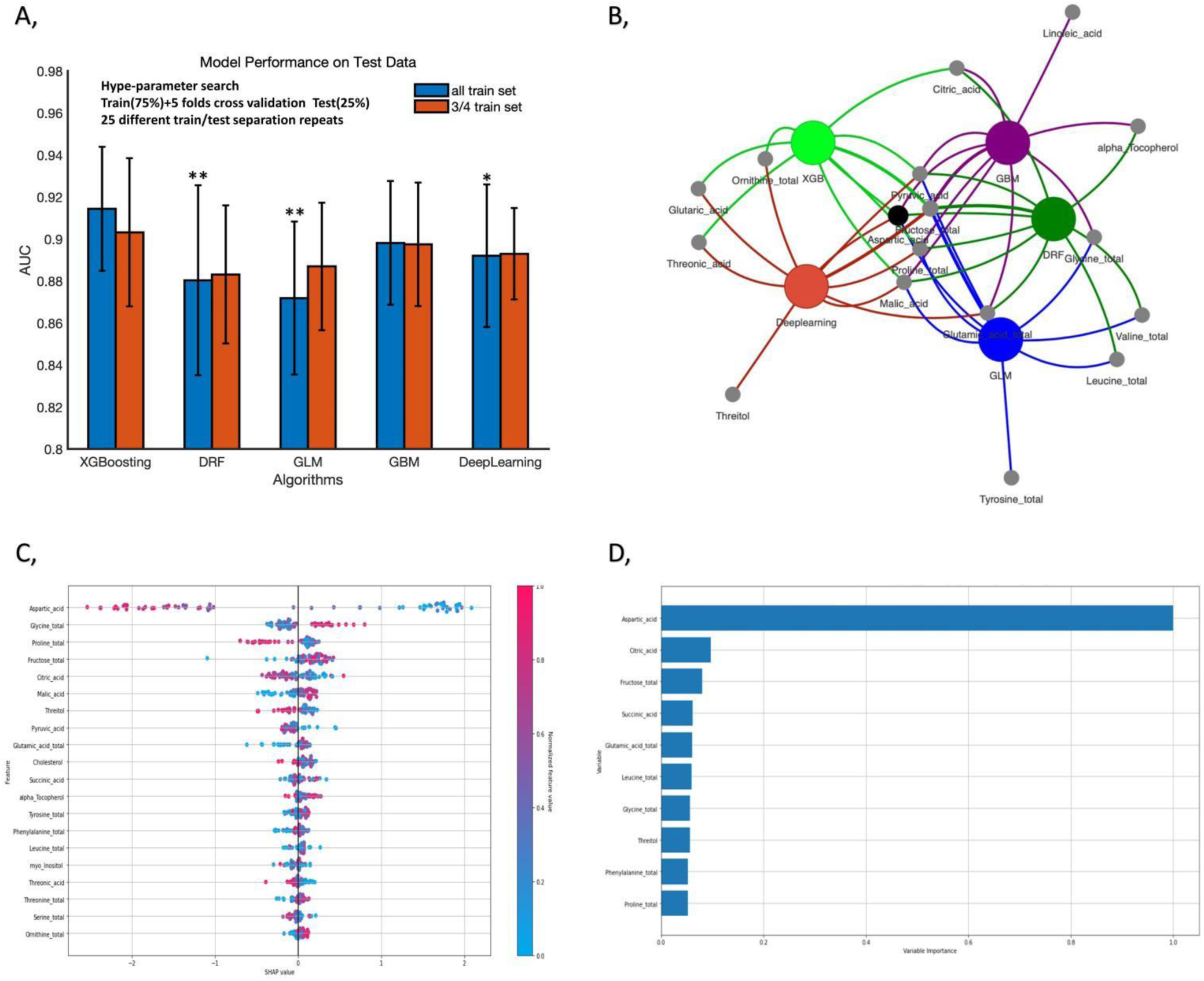
Different machine learning classifiers results. (A) Average AUC on 20 hold out test sets of the XGBoosting algorithm (0.9144) against four other machine learning algorithms for the prediction of two body activity index groups from metabolomics data: Distributed Random Forest and Extremely Randomized Trees (DRF) (0.8804), Generalized Linear Model with regularization (GLM) (0.8719), H2O GBM (GBM) (0.8982) and DeepLearning models (0.8921). For each algorithm, we assess the effect of sample size by building a separated classifier with 1/4 training set removed. (B) Bipartite graph of the top metabolites extracted from the five machine-learning algorithms. For each algorithm, we keep the metabolites if they are identified over 10times in the top ten metabolites over 20 hold out test sets. (C) The variable importance for the XGBoosting algorithm in one hold-out test set. (D) SHAP summary plot of the XGBoosting algorithm in one hold out test set. It shows the contribution of the features for each instance (row of data).

Figure 3A illustrates the averaged AUC values calculated on the 25% hold-out test sets for each algorithm across the 25 random train-test separations. The detailed AUC results are listed in Supplementary material S2. XGBoosting achieved the highest performance with an average AUC value of 0.9150. This result was statistically significant (Wilcoxon signed-rank test P<0.01) compared to the generalized linear models (GLM) that had an average AUC value of 0.8695. The superior performance of XGBoosting suggests the presence of non-linear effects originating from the metabolic network systems. To assess the effect of sample size on various classifiers’ performance, we randomly removed a quarter of the training sets and evaluated the five algorithms. The AUC accuracy of each algorithm hardly changed. Surprisingly, DRF, GLM, and Deep Learning showed improved AUC accuracy with fewer samples. This effect may be attributed to the presence of outlier samples in the original dataset, which introduced noise during the training process, resulting in poorer performance.

In order to assess the importance of metabolites directly related to the two body activity groups, we ranked the metabolites extracted from the five algorithms based on the testing dataset. We identified the top 10 metabolites for each algorithm by calculating the average variable importance across the 25 repeats. The algorithm-metabolite bipartite graph is shown in Figure 3B, where Aspartate, Proline, Fructose, Pyruvate and Malic Acid were consistently identified as the top metabolites across almost all classifiers. The detailed metabolite importance values of each algorithm are presented in Supplementary material S2.

For a better understanding of the variable importance results, we applied a multi-algorithm auto-machine learning approach, including all five algorithms with a maximum of 100 models, using the ‘automl’ function in the H2o.py package. XGBoosting demonstrated the best performance, as shown in Supplementary Table S1. The Pareto front plot in Supplementary Figure S3 determined the optimal subset classifier, which included XGBoosting and GBM classifiers, highlighting the superiority of boosting methods for this task. Figure 3C and 3D present the variable importance and SHAP summary plot for the leading XGBoosting classifier on the test set. The analysis revealed that Aspartate was the most important metabolite, accounting for over 90% of the importance. This highlights the direct influence of the metabolomics aspect on the body activity index. The Spearman’s correlation heatmap shown in Figure 4 further supports this observation, with Aspartate exhibiting the most significant correlation with body strength data. Although other metabolites, such as Proline, Malic Acid, and Pyruvate, had lower importance values, they consistently appeared among the top 10 metabolites across different classifiers. In figure 4, we also did the t-test for all metabolites between the two groups, where the differences with significance are plotted. Interestingly, they didn’t fully match the classifier results, e.g. Pyruvate is identified as key metabolites by all classifiers but didn’t show significance. This may suggest that the effect of Pyruvate is non-linear between two groups. In addition, as shown in Figure 3D, the SHAP plot of the classifier top metabolites still shows good separation between two groups, albeit with less pronounced distinctions compared to Aspartate. This further indicates that they play a role in reflecting non-linear metabolic effects on the body activity index. We choose the eight most important metabolites: Aspartate, Proline, Fructose, Malic Acid, Pyruvate, Valine, Citrate and Ornithine, and map them to the KEGG pathways as shown in Supplementary Figure S4. We can see aside from a few large comprehensive pathways, the top metabolites identified in the classifier results are most related to Central carbon metabolism in cancer and 2-Oxocarboxylic acid metabolism. However, it merely revealed a surface-level connection between active aging and these pathways, which falls short of providing a comprehensive understanding of the underlying biochemical regulations of the active aging dynamics.

**Figure 4.**
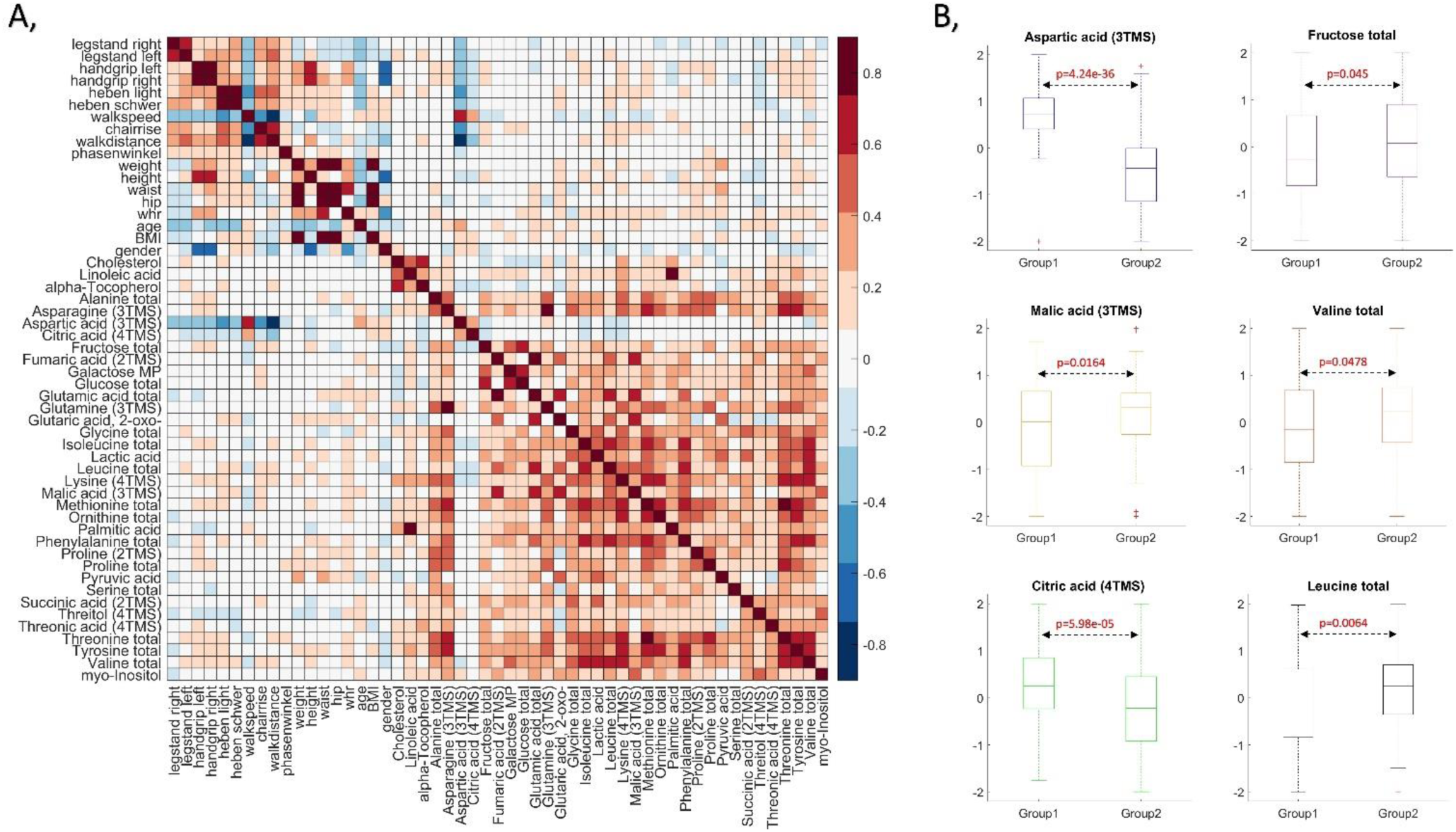
(A) heatmap of the Pearson correlation matrix between the measured physical data and metabolomics data and (B) metabolites’ concentrations comparison between two groups.

### Predictive inverse metabolic interaction modelling using the COVRECON platform

While the machine learning and classifier results provide insights into the variable importance between the measured metabolites and the body activity index, this does not explain the mechanistic change between the two groups. Since for each old adult, metabolomics analysis was done three times, first time point, after 3 months and after 6 months, we plotted the correlation heatmap of the change of all body features and metabolomics measurement changes within two the time intervals in Figure 5. It is evident that the correlation patterns within the metabolomics measurement changes show high similarity. This reflects the internal dynamics of the metabolic networks. Nevertheless, when we check the highly correlated metabolites, we may find no biochemical reactions between the two metabolites from any database. This situation frequently happens, e.g. in Figure 5, Threonine, Tyrosine and Valine show a high correlation, yet no direct biochemical reactions occur among them. This is because the high correlations originate from the network dynamics. Thus, finding the causal interactions among the metabolites is crucial.

**Figure 5.**
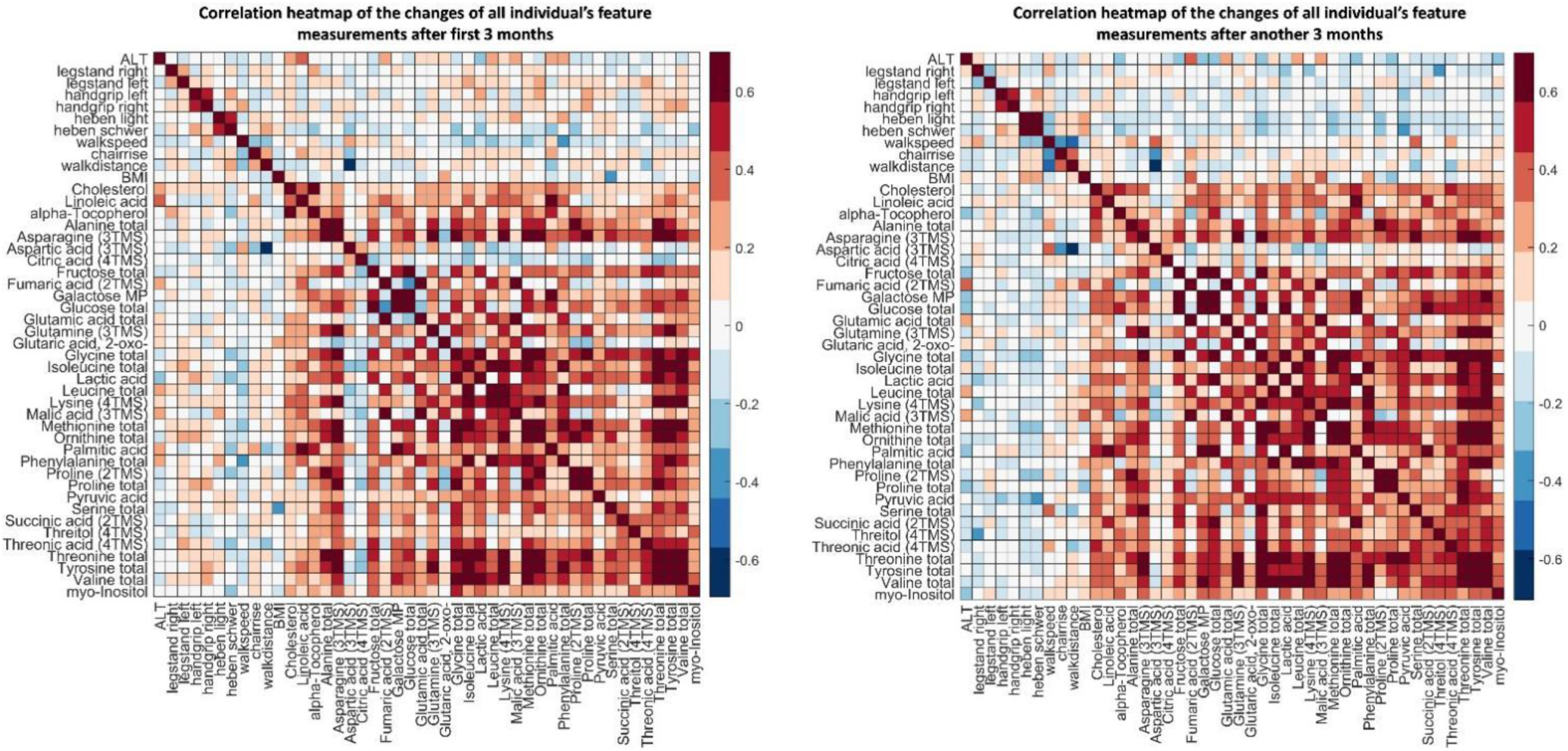
heatmaps of the Pearson correlation matrix of all body features and metabolomics measurement changes within two the time intervals (left: first 3 months, right: another 3 months)

In recent years, inverse differential Jacobian algorithms have been developed, providing a convenient way to infer causal dynamics of metabolic networks from metabolomics data (Nägele, et al., 2014; Sun and Weckwerth, 2012; Weckwerth, 2019; Wilson, et al., 2020; Li, et al., 2023). Besides the metabolomics measurements, metabolic reconstruction is used as complementary information to build a topological model for metabolic interaction network. Based on this, we have developed the COVRECON toolbox (available at: https://bitbucket.org/mosys-univie/covrecon/src/main/) (Li, et al., 2023). As shown in the method part, we applied the COVRECON workflow to the two group datasets. The COVRECON workflow consists of two steps: building the metabolic interaction network and the inverse Jacobian calculation.

As described in COVRECON (Li, et al., 2023), we used a default setting in the Sim-Network part to generate a metabolic super-pathway network of the measured metabolites. Each edge in the network represents a feasible pathway between two nodes (metabolites) and reflects a non-zero component in the system Jacobian matrix. The default setting assigns a fixed weight of one to each reaction, and the reverse reaction weight is based on the log value of its delta Gibbs free energy. Additionally, a pathway-steps limitation of 4 is set. Detailed information about reactions, enzymes and genes of the resulting metabolic interaction network can be found in the Supplemental Material S3. By integrating the covariance of the metabolomics data from both groups and the Jacobian structure matrix, we can perform the inverse Jacobian analysis in the second part of COVRECON toolbox. The COVRECON workflow and toolbox address the ill-conditioned matrix problem associated with the inverse Jacobian approach through a regression loss-based algorithm, significantly improving its stability and feasibility (Li, et al., 2023; Sun, et al., 2015). However, given that the inverse Jacobian approach is based on the Jacobian structure and is more reliable in smaller-sized models, we selected a tailored core part of the whole model containing 10-20 metabolites based on the classifier variable importance results as described in method part. The same network reduction strategy as in Sim-Network was employed, with additional indirect connections added to the reduced model. For example, an additional connection from Proline to Aspartate was added to account for the indirect effects through the connections from Proline to Asparagine and from Asparagine to Aspartate (Figure 6). Figure 5 presents 12 typical results in the repeated calculation. All the repeated results are available in Supplementary material S6. It is evident that even though the local results are different due to the influence from the Jacobian structure information, the Inverse Jacobian approach shows stability on several highlighted metabolic interactions. For example, the interactions Proline->Aspartate, Ornithine->Aspartate, Citrate->Aspartate and Glutamate->2-oxo glutaric acid are high valued in the resulted differential metabolic interaction network of many repeats. To present the overall metabolic interaction importance, we integrated all the 200 local results into the full differential Jacobian (DJ) by calculating the average value of each metabolic interaction within the repeats. The final R* matrix and the differential interaction network is presented in Figure 6A & 6B respectively. In Figure 6B, we plot only the highlighted metabolic interactions with calculated value (scaled to 0-1) above 0.5. Here we note that, the result showed robustness, with similar overall R* using 100, 200 and 500 repeats. Further results are using 200 repeats.

**Figure 6.**
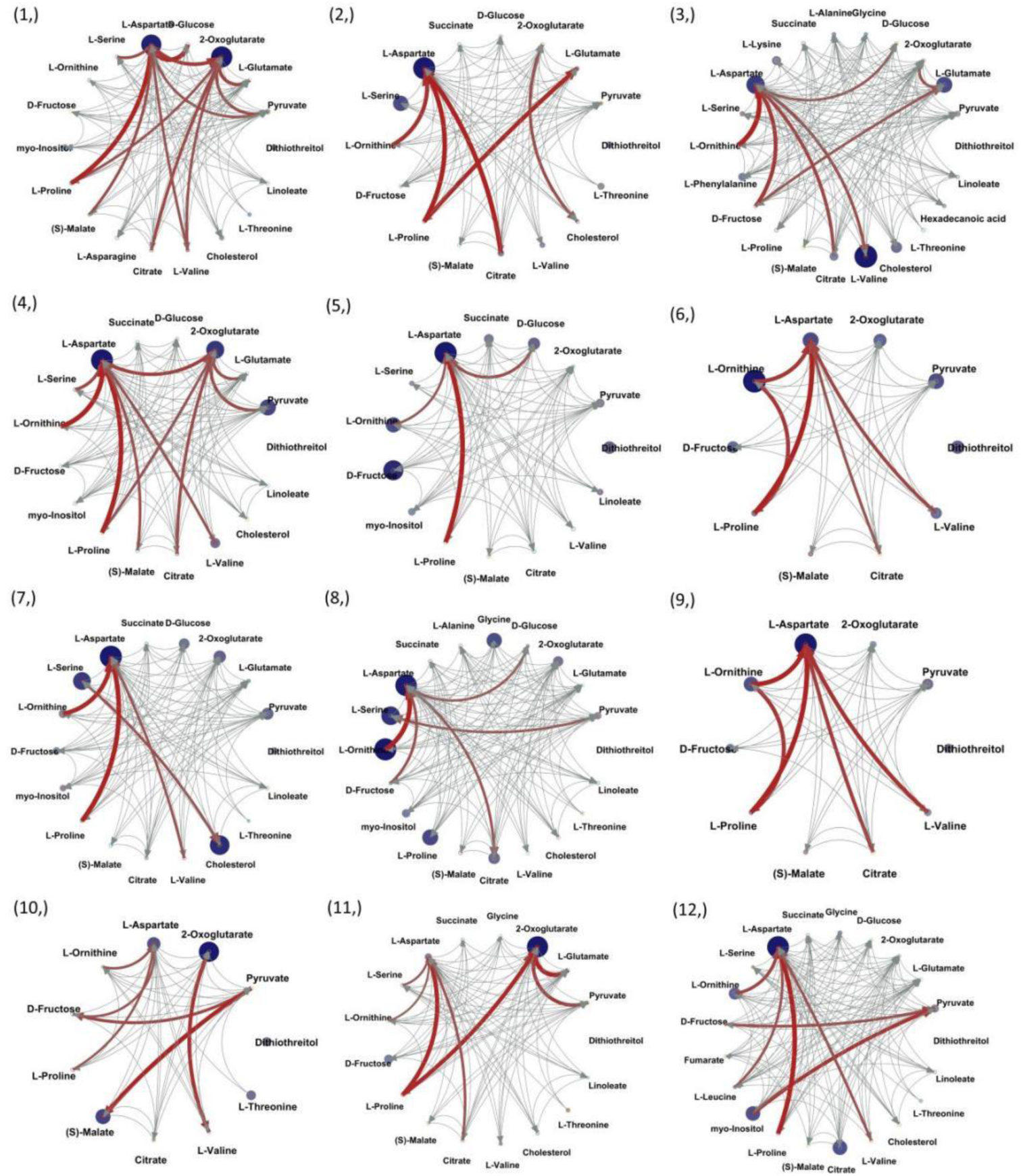
Biochemical pathway network reconstruction and inverse Jacobian calculation repeats with 12 different local Jacobian structures Using COVRECON tool. The detailed information of each metabolic interaction can be interactively checked in the matlab figure format result in Supplementary Material. An excel file listing all interaction information is available in the supplementary material.

Through this COVRECON approach, we are able to find several important perturbed metabolic interactions between the two body activity index clustered groups. The highlighted interactions and the detailed reactions, enzymes and gene information are presented in Supplementary material S4. These findings provide valuable insights into the regulatory interactions and dynamics of the metabolic network related to Aspartate, further supporting its importance as the dominant biomarker in the classifiers results. As shown in Figure 7C, several reactions are consistently identified in several highlighted metabolic interactions. Among these, enzyme aspartate transaminase (AST, EC number 2.6.1.1) is identified in 11 out of the 15 highlighted interactions and shown in all the largest valued interactions: Proline->Aspartate, Valine->Aspartate, Citrate->Aspartate and Glutamate->2-oxo glutaric acid. The enzyme Glutamic-Pyruvic Transaminase (ALT, EC number: 2.6.1.2) is also highlighted. Notably, both AST and ALT are important enzymes in amino acid metabolism, and recently there is indication of their involvement in health related issues of older adults (Goh, et al., 2015; Le Couteur, et al., 2010; Nakajima, et al., 2022). Furthermore, enzyme asparagine synthetase B (EC number: 6.3.5.4) was identified in 8 out of the 15 highlighted interactions. This enzyme is less studied for health issues of elderly peoples. However, asparagine synthetase (ASNS) deficiency was recently discovered as a metabolic disorder of non-essential amino acids (Yamamoto, et al., 2017). Moreover, it is evident that most identified enzymes in Figure 6c belong to enzyme class of transaminases (EC:2.6.1.-). The transaminase enzymes are important in the production of various amino acids, and measuring the concentrations of various transaminases in the blood is important in diagnosing and tracking of many diseases (Oh, et al., 2017).

**Figure 7.**
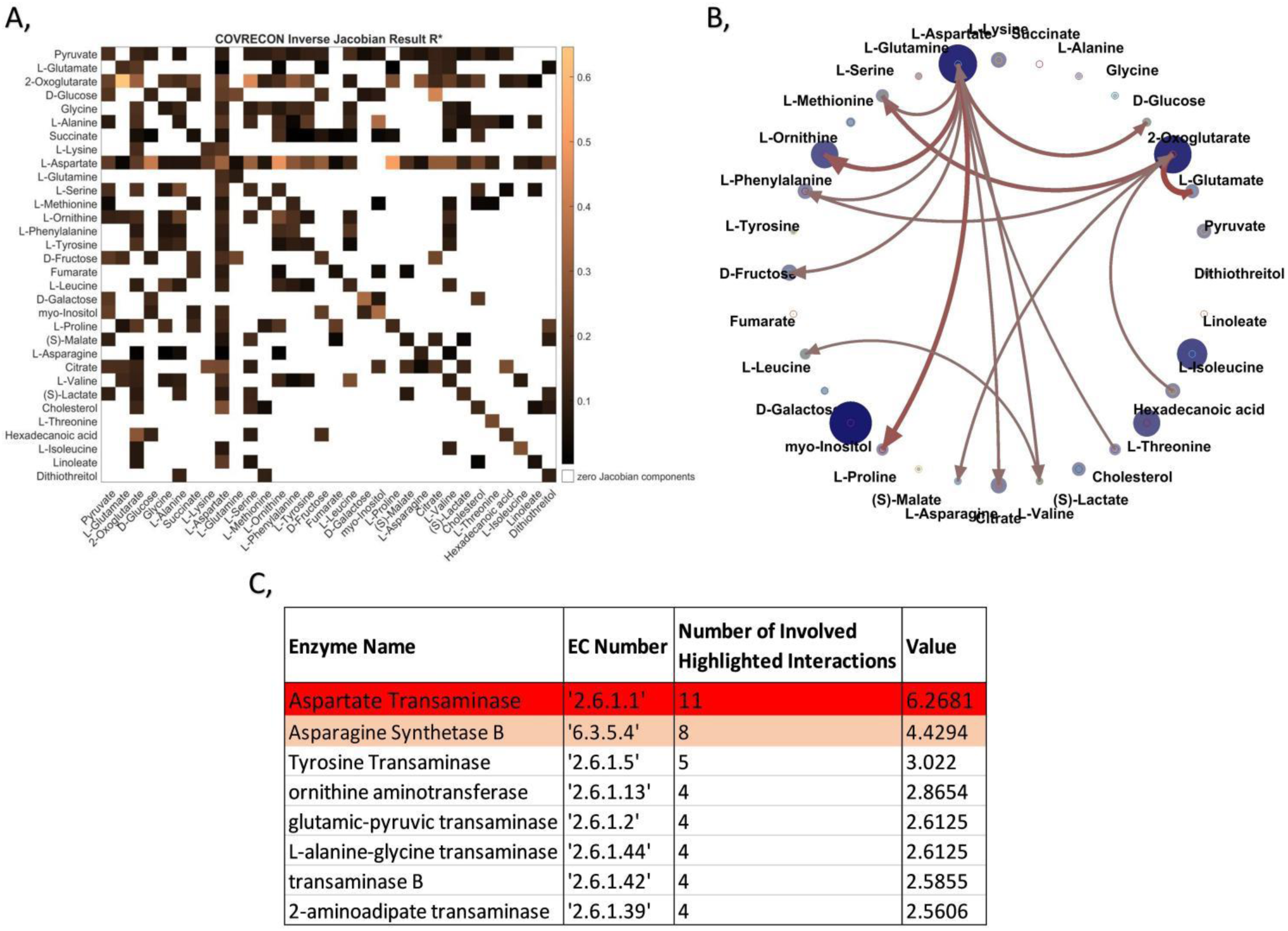
The overall inverse Jacobian results integrating all 200 local calculation repeats. Figure 7A is the average inverse Jacobian matrix; Figure 7B only plotted highlighted metabolic interactions with inverse Jacobian calculation value above 0.5 (scaled to 0-1); Figure 7C listed the most relevant enzymes, where Aspartate Transaminase showed 11 times out of the 15 highlighted interactions, with the accumulated value 6.2681.

For a further analysis of the enzymes, we conducted routine blood tests measurements of the old adults across the three time points. Four metabolic enzymes were measured: AST, ALT, Gamma-glutamyltransferase (GGT) and Creatine Kinase (CK). The data measurements are presented in Supplementary material S2. As shown in Figure 8 and Supplementary Figure S6, we compared the enzyme measurements between the two groups (active/less active). The results suggested significant differences in AST and ALT, while GGT and CK did not exhibit such significant variations. This observation validates the inverse Jacobian results in Figure 7. Furthermore, we compared the AST and ALT changes within the two 3-months’ time intervals. As demonstrated in Figure 7, both AST and ALT showed significant changes in the “active group”, while the changes were not significant in the “less active group” during both 3-months intervals. Notably, the changes also exhibited significant differences between the two groups. Specifically, in the “active group”, AST and ALT demonstrated a significant larger decrease during the first 3 months, followed by a significant larger increase in the subsequent 3-months interval. This suggests that a larger plasticity of enzymatic liver and muscle systems in individuals with a high level of body activity. Interestingly, a few studies have revealed similar observations while investigating the enzyme variations. In a long-term study of 29 routine laboratory measurements of 30 athletes, AST and ALT exhibited significantly larger variations over an 11-months period compared to those reported for general population (Diaz-Garzon, et al., 2022; Diaz-Garzon, et al., 2023). Moreover, various studies have evidenced the enzyme fluctuations within healthy individuals’ blood samples from physical activity and exercises (Nunez, et al., 2019; Pavletic and Wright, 2015; Pettersson, et al., 2008; Ruiz, et al., 2014; Taylor, 2022; Tiller and Stringer, 2023).

**Figure 8.**
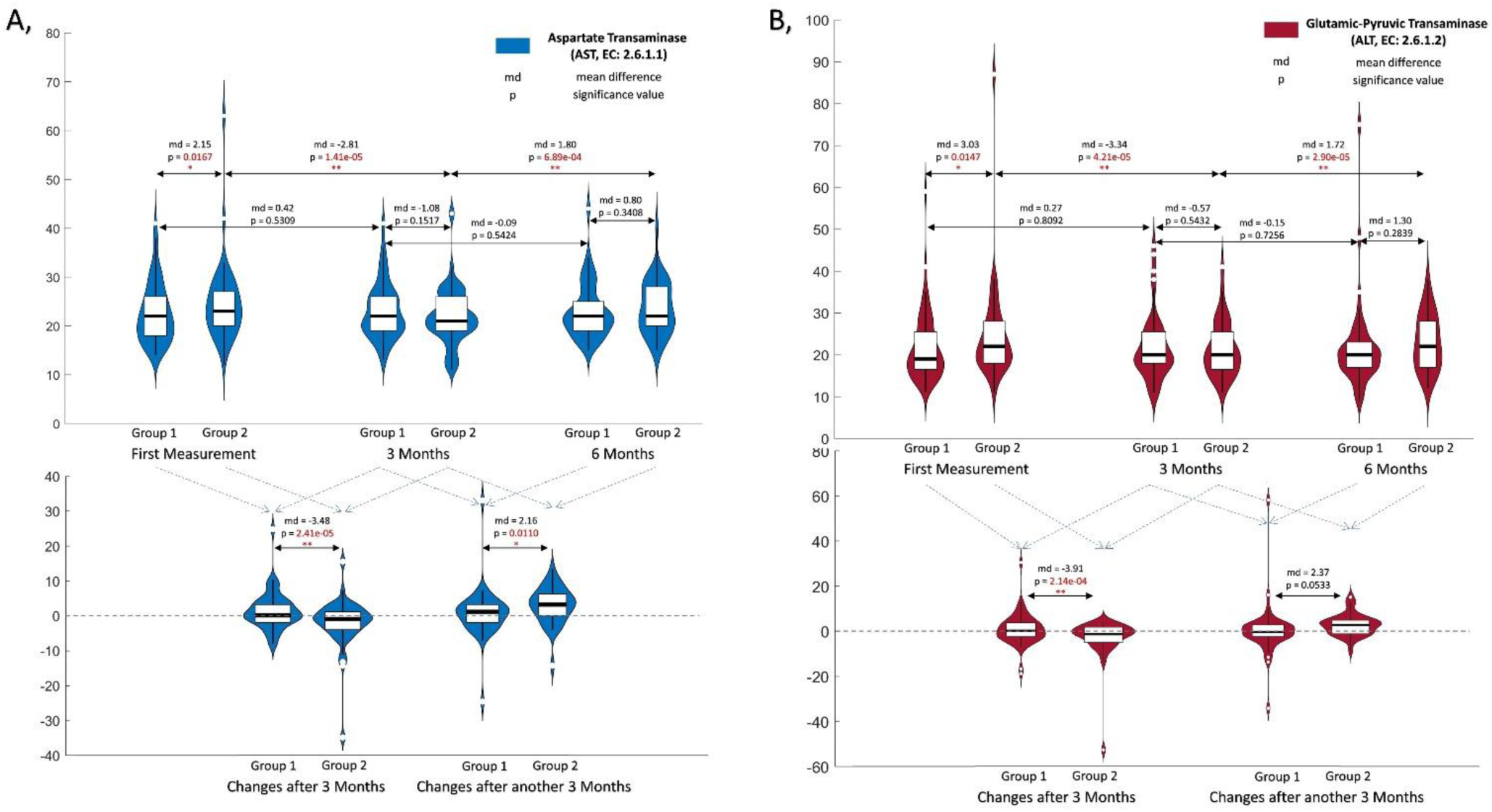
The enzyme measurements for the two enzymes identified in inverse Jacobian results: (A) Aspartate Transaminase (AST) and (B) Glutamic-Pyruvic Transaminase (ALT) The enzymes are measured in three time points: first measurement, 3 months later and 6 months later. The enzymes’ concentrations are compared between two groups (physical active and less active groups) and different time points, where significances are highlighted.

## Discussion

In this article, we measured 263 plasma metabolomics samples to study active aging and fitness in a cohort of very old adults close to or above the average life expectancy. Using a CCA approach, we clustered all old adults and samples into two groups based on a body activity index. Then we identified several key biomarkers between these two groups through machine- and deep learning analysis. The identified metabolites are Aspartate, Proline, Fructose, Malic Acid, Pyruvate, Valine, Citrate and Ornithine, where Aspartate showed dominant effects. XGboosting showed the best performance. In a further analysis, we applied the COVRECON (Li, et al., 2023) approach to the two group metabolomics datasets. Through this method, we identified several key metabolic interaction changes between the two active-less active groups. Many of these interactions are related to aspartate, this is consistent with the machine learning results. By checking the detailed enzyme information of the highlighted metabolic interactions, we identified several important enzyme regulations. The enzyme AST showed a relation to most highlighted interactions. The blood measurements of all individuals across the three time points validate the results. Existing studies also showed that AST and ALT is highly related to health issues of older adults (Andy and Keeffe, 2003).

### Metabolomics chances for resistance training

As shown in Supplementary table S2, we conducted a group difference t-test for the metabolomics measurements. Where Alpha Tocopherol shows significant difference between nutritional supplement intake group (E) and the other two groups, as it is a part of the supplement FortiFit. The metabolites Linnileic acid, Methionine, Palmitic acid, Succinate and Tyrosine show a significant difference between the control group (K) and the resistance training groups (T & E). Interestingly, this divergence contrasts with the results obtained from the body activity classifiers, suggesting distinct metabolic mechanisms for resistance exercise and endurance exercise. This mechanistic difference between endurance and resistance exercise has been previously explored (Morville, et al., 2020), where the metabolites changes induced by endurance or resistance exercise are identified in two different modes.

Moreover, several studies have reported that endurance exercise but not resistance exercise has a high relevance to aging related questions. In Cao Dinh, et al., 2019, among 100 old women (aged over 65 years) the study reported that strength endurance training significantly reduces senescence-prone T cells, which is widely recognized as age-related (Childs, et al., 2015),while intensive training showed no significant influence. In another study, Weiner, et al., 2019 concluded that endurance but not resistance training has anti-aging effects while examining a total of 124 healthy previously inactive individuals (Werner, et al., 2019). These studies provide additional support for our body activity index and metabolic network analysis.

### Aspartate as a blood biomarker for body activity

Aspartic acid is one of the 22 protein-generic amino acids. It is involved in the malate-aspartate shuttle, which facilitates the transfer of electrons and energy between the cytoplasm and mitochondria, ultimately contributing to the production of ATP and the efficient functioning of cellular energy metabolism (Borst, 2020). Thus, it is particularly important in tissues with high-energy demands, such as muscle, liver and the heart. This may account for the larger aspartate metabolism in the “active group”. From this point, several groups have evidenced the effect of aspartate as an important supplement for attenuation of exercise-induced hyperammonaemia and an increase in exercise endurance (Marquezi, et al., 2003; Trudeau, 2008). On the other hand, aspartate is involved in the removal of ammonia from the body through the urea cycle (Fibriansah, et al., 2011). Performing exercise can lead to ammonia production as a byproduct of energy metabolism. Aspartate may be used to help detoxifying ammonia, potentially altering its levels.

### Old adults with better body activity have larger plasticity of enzymatic liver and muscle system

AST and ALT are two of the routine blood test enzymes highly related to individual’s liver but also muscle and heart health (Lala, et al., 2022), where elevated levels of AST and ALT enzymes beyond a specified threshold may indicate medical condition like hepatitis, liver disease or myonecrosis. The ratio AST/ALT is a significant sign of liver disease. We plotted the AST/ALT ratio changes over the three time points for the two groups in Supplementary Figure S7. The results showed no significant changes across the time points and groups. This suggests that AST and ALT variations originates from non-disease related factors. Furthermore, scientific investigations have furnished evidence supporting the notion that physical exercise and improved fitness levels can also lead to a transient elevation of these enzyme levels within a healthy range for individuals without underlying liver issues (Pavletic and Wright, 2015; Pettersson, et al., 2008; Tiller and Stringer, 2023). This exercise-induced transaminase elevation is a well-documented phenomenon, commonly observed in response to vigorous physical activity. It is essential to recognize that these exercise-related increases in AST and ALT levels are typically temporary and return to baseline levels shortly after physical exertion. This indicates larger AST and ALT variations for individuals with better body functionality/activity, as observed in Figure 8. This viewpoint is also suggested in a long-term study of 29 routine laboratory measurements of 30 athletes, where AST and ALT exhibited significantly larger variations over an 11-months period compared to those reported for general population (Diaz-Garzon, et al., 2022; Diaz-Garzon, et al., 2023).

In conclusion, this study is the first time we integrate machine learning statistical analysis and COVRECON inverse Jacobian analysis together. In metabolomics analysis, machine learning based statistical methods aids us find the key metabolites. As for the dynamical analysis, aside from kinetic modeling which needs a large number of parameter fitting processes, we showed the predictive metabolic interaction modelling using the inverse differential Jacobian approach. This novel approach might be highly relevant to find the important dynamic regulations between two conditions. By integrating the machine learning results, we showed a robust approach for the inverse differential Jacobian calculation.

## References

Akbari, A., Haiman, Z.B. and Palsson, B.O. A data-driven approach for timescale decomposition of biochemical reaction networks. Msystems 2024;9(2):e01001-01023.

Alakwaa, F.M., Chaudhary, K. and Garmire, L.X. Deep learning accurately predicts estrogen receptor status in breast cancer metabolomics data. J Proteome Res 2018;17(1):337–347.

Andy, S.Y. and Keeffe, E.B. Elevated AST or ALT to nonalcoholic fatty liver disease: accurate predictor of disease prevalence? In.: LWW; 2003. p. 955–956.

Balashova, E.E., et al. Metabolome Profiling in Aging Studies. Biology 2022;11(11):1570.

Borst, P. The malate–aspartate shuttle (Borst cycle): How it started and developed into a major metabolic pathway. Iubmb Life 2020;72(11):2241–2259.

Boudiny, K. and Mortelmans, D. A critical perspective: towards a broader understanding of’active ageing’. E-journal of Applied Psychology 2011;7(1):8–14.

Bruzzone, C., et al. Metabolomics as a powerful tool for diagnostic, pronostic and drug intervention analysis in COVID-19. Frontiers in Molecular Biosciences 2023;10:1111482.

Cao Dinh, H., et al. Strength endurance training but not intensive strength training reduces senescence-prone T cells in peripheral blood in community-dwelling elderly women. The Journals of Gerontology: Series A 2019;74(12):1870–1878.

Caprara, M., et al. Active aging promotion: results from the Vital Aging Program. Current Gerontology and Geriatrics Research 2013;2013.

Chen, T. and Guestrin, C. Xgboost: A scalable tree boosting system. In, Proceedings of the 22nd acm sigkdd international conference on knowledge discovery and data mining. 2016. p. 785–794.

Childs, B.G., et al. Cellular senescence in aging and age-related disease: from mechanisms to therapy. Nature medicine 2015;21(12):1424–1435.

Diaz-Garzon, J., et al. Long-term within-and between-subject biological variation of 29 routine laboratory measurands in athletes. Clinical Chemistry and Laboratory Medicine (CCLM) 2022;60(4):618–628.

Diaz-Garzon, J., et al. Long-Term Within-and Between-Subject Biological Variation Data of Hematological Parameters in Recreational Endurance Athletes. Clinical Chemistry 2023;69(5):500–509.

Dunn, W.B., et al. Procedures for large-scale metabolic profiling of serum and plasma using gas chromatography and liquid chromatography coupled to mass spectrometry. Nature protocols 2011;6(7):1060–1083.

Fernández-Ballesteros, R., et al. Active aging: a global goal. In.: Hindawi; 2013.

Feurer, M. and Hutter, F. Hyperparameter optimization. In, Automated machine learning. Springer, Cham; 2019. p. 3-33.

Fibriansah, G., et al. Structural basis for the catalytic mechanism of aspartate ammonia lyase. Biochemistry 2011;50(27):6053–6062.

Ghini, V., et al. Serum NMR profiling reveals differential alterations in the lipoproteome induced by pfizer-BioNTech vaccine in COVID-19 recovered subjects and naïve subjects. Frontiers in molecular biosciences 2022;9:839809.

Goh, G.B.-B., et al. Age impacts ability of aspartate–alanine aminotransferase ratio to predict advanced fibrosis in nonalcoholic fatty liver disease. Digestive diseases and sciences 2015;60:1825–1831.

Gonzalez-Covarrubias, V., Martínez-Martínez, E. and del Bosque-Plata, L. The potential of metabolomics in biomedical applications. Metabolites 2022;12(2):194.

Haiman, Z.B., et al. MASSpy: Building, simulating, and visualizing dynamic biological models in Python using mass action kinetics. PLoS computational biology 2021;17(1):e1008208.

Hardoon, D.R., Szedmak, S. and Shawe-Taylor, J. Canonical correlation analysis: An overview with application to learning methods. Neural computation 2004;16(12):2639–2664.

Havighurst, R.J. Successful aging. Processes of aging: Social and psychological perspectives 1963;1:299-320.

Jamshidi, N. and Palsson, B.Ø. Mass action stoichiometric simulation models: incorporating kinetics and regulation into stoichiometric models. Biophysical journal 2010;98(2):175–185.

Ke, G., et al. Lightgbm: A highly efficient gradient boosting decision tree. Advances in neural information processing systems 2017;30.

King, Z.A., et al. BiGG Models: A platform for integrating, standardizing and sharing genome-scale models. Nucleic acids research 2016;44(D1):D515–D522.

Kohl, H.W., et al. The pandemic of physical inactivity: global action for public health. The lancet 2012;380(9838):294-305.

Kügler, P. and Yang, W. Identification of alterations in the Jacobian of biochemical reaction networks from steady state covariance data at two conditions. Journal of Mathematical Biology 2014;68(7):1757–1783.

Lala, V., Zubair, M. and Minter, D.A. Liver function tests. In, StatPearls [internet]. StatPearls Publishing; 2022.

Le Couteur, D.G., et al. The association of alanine transaminase with aging, frailty, and mortality. Journals of Gerontology Series A: Biomedical Sciences and Medical Sciences 2010;65(7):712–717.

LeCun, Y., Bengio, Y. and Hinton, G. Deep learning. nature 2015;521(7553):436-444.

Li, J., Waldherr, S. and Weckwerth, W. COVRECON: Automated Integration of Genome- and Metabolome-Scale Network Reconstruction and Data-driven Inverse Modeling of Metabolic Interaction Neworks. Bioinformatics 2023.

Li, J., Waldherr, S. and Weckwerth, W. COVRECON: automated integration of genome- and metabolome-scale network reconstruction and data-driven inverse modeling of metabolic interaction networks. Bioinformatics 2023;39(7).

Li, J., Waldherr, S. and Weckwerth, W. COVRECON: automated integration of genome-and metabolome-scale network reconstruction and data-driven inverse modeling of metabolic interaction networks. Bioinformatics 2023;39(7):btad397.

Liebal, U.W., et al. Machine learning applications for mass spectrometry-based metabolomics. Metabolites 2020;10(6):243.

Malkowski, O.S., Kanabar, R. and Western, M.J. Socio-economic status and trajectories of a novel multidimensional metric of Active and Healthy Ageing: the English Longitudinal Study of Ageing. Scientific Reports 2023;13(1):6107.

Marquezi, M.L., et al. Effect of aspartate and asparagine supplementation on fatigue determinants in intense exercise. International journal of sport nutrition and exercise metabolism 2003;13(1):65–75.

Meoni, G., et al. Metabolomic/lipidomic profiling of COVID-19 and individual response to tocilizumab. PLoS Pathogens 2021;17(2):e1009243.

Morville, T., et al. Plasma metabolome profiling of resistance exercise and endurance exercise in humans. Cell reports 2020;33(13).

Nägele, T. Metabolic regulation of subcellular sucrose cleavage inferred from quantitative analysis of metabolic functions. Quantitative Plant Biology 2022;3:e10.

Nägele, T., et al. Solving the differential biochemical Jacobian from metabolomics covariance data. PloS one 2014;9(4):e92299.

Nakajima, K., et al. High aspartate Aminotransferase/Alanine aminotransferase ratio may be Associated with all-cause mortality in the Elderly: a Retrospective Cohort Study using Artificial Intelligence and Conventional Analysis. In, Healthcare. MDPI; 2022. p. 674.

Nunez, D.J., et al. Factors influencing longitudinal changes of circulating liver enzyme concentrations in subjects randomized to placebo in four clinical trials. American Journal of Physiology-Gastrointestinal and Liver Physiology 2019;316(3):G372–G386.

Oesen, S., et al. Effects of elastic band resistance training and nutritional supplementation on physical performance of institutionalised elderly—A randomized controlled trial. Experimental gerontology 2015;72:99–108.

Offerman, J., et al. Attitudes related to technology for active and healthy aging in a national multigenerational survey. Nature Aging 2023;3(5):617–625.

Oh, R.C., et al. Mildly elevated liver transaminase levels: causes and evaluation. American family physician 2017;96(11):709–715.

Panyard, D.J., Yu, B. and Snyder, M.P. The metabolomics of human aging: Advances, challenges, and opportunities. Science Advances 2022;8(42):eadd6155.

Patti, G.J., Yanes, O. and Siuzdak, G. Metabolomics: the apogee of the omics trilogy. Nature reviews Molecular cell biology 2012;13(4):263–269.

Pavletic, A.J. and Wright, M.E. Exercise-induced elevation of liver enzymes in a healthy female research volunteer. Psychosomatics 2015;56(5):604.

Pettersson, J., et al. Muscular exercise can cause highly pathological liver function tests in healthy men. British journal of clinical pharmacology 2008;65(2):253–259.

Pomyen, Y., et al. Deep metabolome: Applications of deep learning in metabolomics. Computational and Structural Biotechnology Journal 2020;18:2818–2825.

Ruiz, J.R., et al. Physical activity, sedentary time, and liver enzymes in adolescents: the HELENA study. Pediatr. Res. 2014;75(6):798–802.

Sidak, D., et al. Interpretable machine learning methods for predictions in systems biology from omics data. Frontiers in Molecular Biosciences 2022;9:926623.

Sindelar, M., et al. Longitudinal metabolomics of human plasma reveals prognostic markers of COVID-19 disease severity. Cell Reports Medicine 2021;2(8).

Steuer, R., et al. Structural kinetic modeling of metabolic networks. Proceedings of the National Academy of Sciences 2006;103(32):11868–11873.

Steuer, R., et al. Observing and interpreting correlations in metabolomic networks. Bioinformatics 2003;19(8):1019–1026.

Su, Y., et al. Multi-omics resolves a sharp disease-state shift between mild and moderate COVID-19. Cell 2020;183(6):1479–1495. e1420.

Sun, X., Länger, B. and Weckwerth, W. Challenges of inversely estimating jacobian from metabolomics data. Frontiers in bioengineering and biotechnology 2015;3:188.

Sun, X. and Weckwerth, W. COVAIN: a toolbox for uni-and multivariate statistics, time-series and correlation network analysis and inverse estimation of the differential Jacobian from metabolomics covariance data. Metabolomics 2012;8(1):81–93.

Taylor, A.W. Physiology of exercise and healthy aging. Human Kinetics; 2022.

Teahan, O., et al. Impact of analytical bias in metabonomic studies of human blood serum and plasma. Analytical chemistry 2006;78(13):4307–4318.

Tiller, N.B. and Stringer, W.W. Exercise-induced increases in “liver function tests” in a healthy adult male: Is there a knowledge gap in primary care? Journal of Family Medicine and Primary Care 2023;12(1):177.

Trudeau, F. Aspartate as an ergogenic supplement. Sports Medicine 2008;38:9–16.

Weckwerth, W. Metabolomics in systems biology. Annual review of plant biology 2003;54(1):669–689.

Weckwerth, W. Metabolomics: an integral technique in systems biology. Bioanalysis 2010;2(4):829–836.

Weckwerth, W. Green systems biology—from single genomes, proteomes and metabolomes to ecosystems research and biotechnology. Journal of proteomics 2011;75(1):284–305.

Weckwerth, W. Unpredictability of metabolism--the key role of metabolomics science in combination with next-generation genome sequencing. Anal Bioanal Chem 2011;400(7):1967–1978.

Weckwerth, W. Unpredictability of metabolism—the key role of metabolomics science in combination with next-generation genome sequencing. Analytical and Bioanalytical Chemistry 2011;400(7):1967–1978.

Weckwerth, W. Toward a unification of system-theoretical principles in biology and ecology—the stochastic lyapunov matrix equation and its inverse application. Frontiers in Applied Mathematics and Statistics 2019;5:29.

Weckwerth, W., Wenzel, K. and Fiehn, O. Process for the integrated extraction, identification and quantification of metabolites, proteins and RNA to reveal their co - regulation in biochemical networks. Proteomics 2004;4(1):78–83.

Weiszmann, J., et al. Metabolome plasticity in 241 Arabidopsis thaliana accessions reveals evolutionary cold adaptation processes. Plant Physiology 2023:kiad298.

Werner, C.M., et al. Differential effects of endurance, interval, and resistance training on telomerase activity and telomere length in a randomized, controlled study. European heart journal 2019;40(1):34–46.

WHO. Active ageing: A policy framework. In.: World Health Organization; 2002.

Wienkoop, S., et al. Integration of metabolomic and proteomic phenotypes: analysis of data covariance dissects starch and RFO metabolism from low and high temperature compensation response in Arabidopsis thaliana. Molecular & Cellular Proteomics 2008;7(9):1725–1736.

Wilson, J.L., et al. Inverse data-driven modeling and multiomics analysis reveals phgdh as a metabolic checkpoint of macrophage polarization and proliferation. Cell Reports 2020;30(5):1542–1552. e1547.

Wongsala, M., Anbäcken, E.-M. and Rosendahl, S. Active ageing–perspectives on health, participation, and security among older adults in northeastern Thailand–a qualitative study. BMC geriatrics 2021;21:1–10.

Yamamoto, T., et al. The first report of Japanese patients with asparagine synthetase deficiency. Brain and Development 2017;39(3):236–242.

Zhou, Z.-H. Ensemble methods: foundations and algorithms. CRC press; 2012.

