## Supplementary_material_S1 for "Machine learning and data-driven inverse modeling of metabolomics unveil key process of active aging"

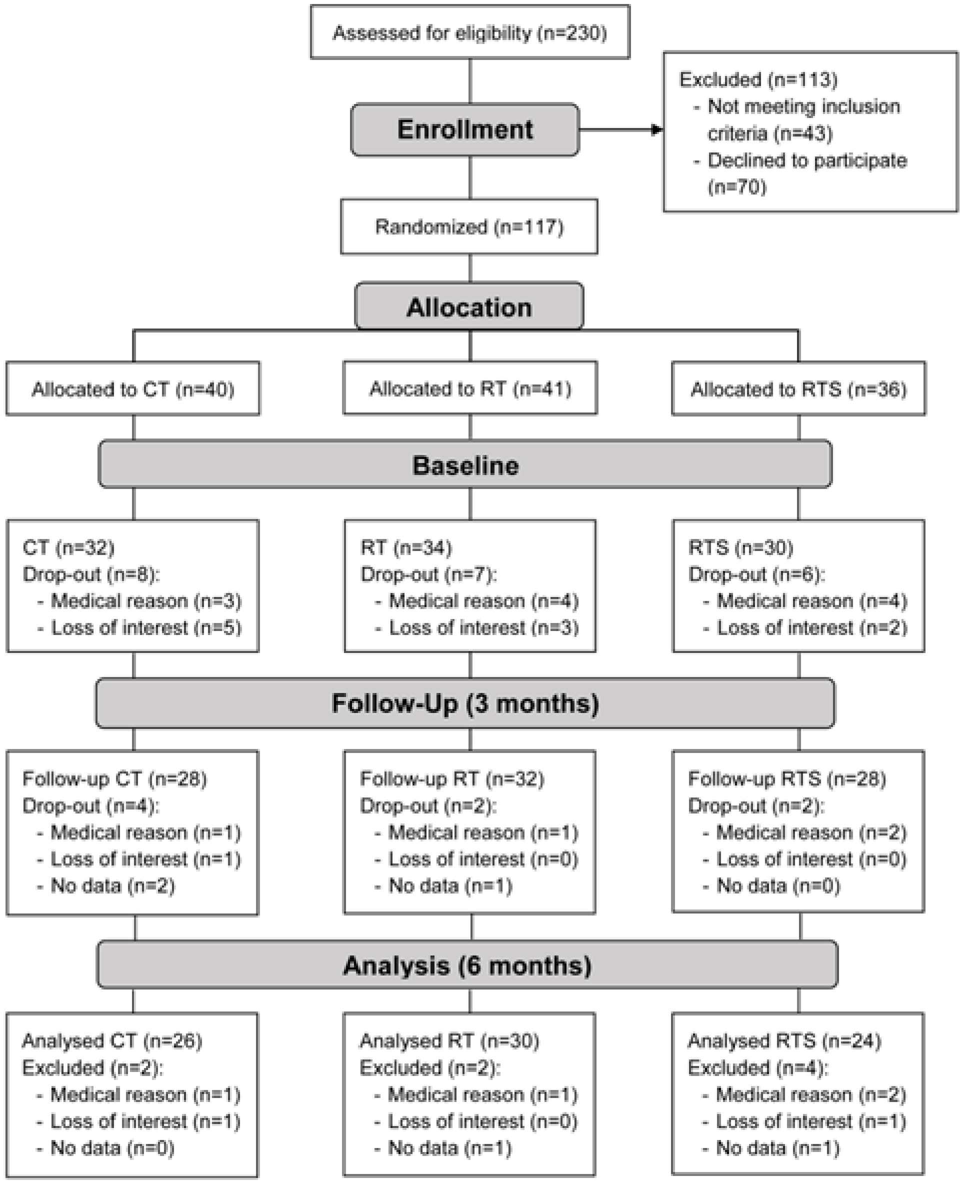


**Supplementary Figure S1 shows experimental study design**

### Treatment

#### Resistance training

The participants performed resistance training twice a week, supervised by a sport scientist. They were able to use elastic bands, chairs and their own body weights. The session consisted of a 10 min warm-up, 30-40 min of strength training that consisted of ten exercises for the main muscle groups (shoulders, arms legs, back, abdomen and chest) and ended with a 10 minutes cool down (Oesen, et al., 2015).

The exertion was adjusted to the participants' individual fitness level by adapting the resistance of the elastic band. 15 repetitions were performed and as soon as the exercise could be easily performed by the subjects, the resistance was increased to perform a more difficult version of the exercise and thus obtain a higher training effect (Oesen, et al., 2015).

#### Resistance training and supplementation

The participants performed the same resistance training as the resistance training group. In addition, they received supplements every day and after each training session. The intake of this supplement was controlled.

Nutritional Supplement FortiFit, produced by NUTRICIA GmbH, Vienna, Austria, contained 20.7g protein (56 energy (En)%, 19.7g whey protein, 3.0g leucine, > 10g essential amino acids), 9.3g carbohydrates (25 En%, 0.8 BE), 3.0g fat (18 En%), 1.2g roughage (2 En%), 800IU (20µg) of vitamin D, 250mg calcium, vitamins C, E, B6 and B12, folic acid and magnesium (Oesen et al. 2015).

#### Cognitive Training

The participants performed twice a week memory training and finger dexterity exercises in sitting position. Therefore, minimizing the "bias" being alone and not being a part in group activities (socialization factor) (Oesen, et al., 2015). Participants of all groups were instructed to maintain their regular food intake.

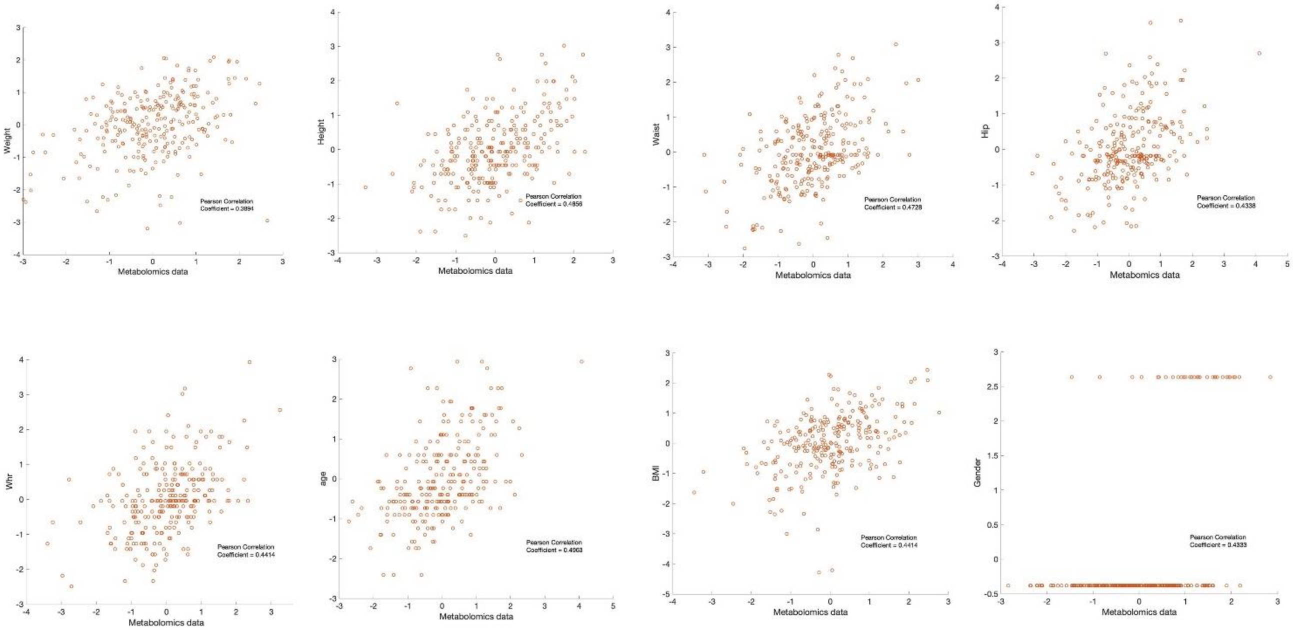


**Figure S2, Canonical analysis of metabolomics data and body size measurements.**

**Table S1, Top 20 models in a 100 maximum models 'automl' result**

| model_id | auc | logloss | aucpr | mean_per_class_error | rmse | mse | training_time_ms |
|---|---|---|---|---|---|---|---|
| XGBoost_grid_1_AutoML_18_20221215_135819_model_5 | 0.960961 | 0.263797 | 0.978493 | 0.0640641 | 0.275694 | 0.0760073 | 368 |
| GBM_grid_1_AutoML_18_20221215_135819_model_11 | 0.954955 | 0.309706 | 0.970467 | 0.106106 | 0.305647 | 0.0934199 | 88 |
| XGBoost_grid_1_AutoML_18_20221215_135819_model_8 | 0.953954 | 0.287761 | 0.96649 | 0.0860861 | 0.296339 | 0.0878166 | 351 |
| XGBoost_grid_1_AutoML_18_20221215_135819_model_7 | 0.948949 | 0.315144 | 0.967971 | 0.11962 | 0.312296 | 0.097529 | 366 |
| XGBoost_2_AutoML_18_20221215_135819 | 0.947948 | 0.301032 | 0.968592 | 0.11962 | 0.300847 | 0.0905089 | 445 |
| GBM_3_AutoML_18_20221215_135819 | 0.947948 | 0.29179 | 0.969714 | 0.0960961 | 0.294029 | 0.0864529 | 120 |
| GBM_grid_1_AutoML_18_20221215_135819_model_2 | 0.946947 | 0.300804 | 0.968795 | 0.0860861 | 0.305624 | 0.0934063 | 146 |
| StackedEnsemble_BestOfFamily_1_AutoML_18_20221215_135819 | 0.943944 | 0.313579 | 0.961715 | 0.0825826 | 0.291704 | 0.085091 | 24499 |
| XGBoost_grid_1_AutoML_18_20221215_135819_model_6 | 0.941942 | 0.310443 | 0.965722 | 0.104605 | 0.312564 | 0.0976965 | 384 |
| GBM_grid_1_AutoML_18_20221215_135819_model_9 | 0.941942 | 0.316609 | 0.964882 | 0.104605 | 0.309971 | 0.0960821 | 103 |
| XGBoost_3_AutoML_18_20221215_135819 | 0.93994 | 0.306946 | 0.964095 | 0.114615 | 0.29881 | 0.0892876 | 345 |
| XGBoost_grid_1_AutoML_18_20221215_135819_model_3 | 0.937437 | 0.357516 | 0.958981 | 0.114615 | 0.328424 | 0.107863 | 370 |
| XGBoost_grid_1_AutoML_18_20221215_135819_model_10 | 0.935936 | 0.337679 | 0.956265 | 0.114615 | 0.323489 | 0.104645 | 356 |
| DeepLearning_grid_1_AutoML_18_20221215_135819_model_3 | 0.934935 | 0.687702 | 0.919111 | 0.0875876 | 0.31974 | 0.102234 | 2610 |
| StackedEnsemble_AllModels_1_AutoML_18_20221215_135819 | 0.934935 | 0.322924 | 0.951681 | 0.0910911 | 0.297026 | 0.0882245 | 32979 |
| XGBoost_grid_1_AutoML_18_20221215_135819_model_4 | 0.933934 | 0.32437 | 0.959287 | 0.104605 | 0.315342 | 0.0994406 | 349 |
| DeepLearning_grid_2_AutoML_18_20221215_135819_model_2 | 0.931431 | 1.68747 | 0.931677 | 0.124625 | 0.366589 | 0.134387 | 4723 |
| DeepLearning_grid_3_AutoML_18_20221215_135819_model_3 | 0.930931 | 1.13271 | 0.936983 | 0.101101 | 0.323924 | 0.104927 | 3992 |
| GBM_2_AutoML_18_20221215_135819 | 0.92993 | 0.31436 | 0.948552 | 0.104605 | 0.306929 | 0.0942057 | 108 |
| GLM_1_AutoML_18_20221215_135819 | 0.926927 | 0.358139 | 0.916452 | 0.0960961 | 0.308512 | 0.0951796 | 80 |

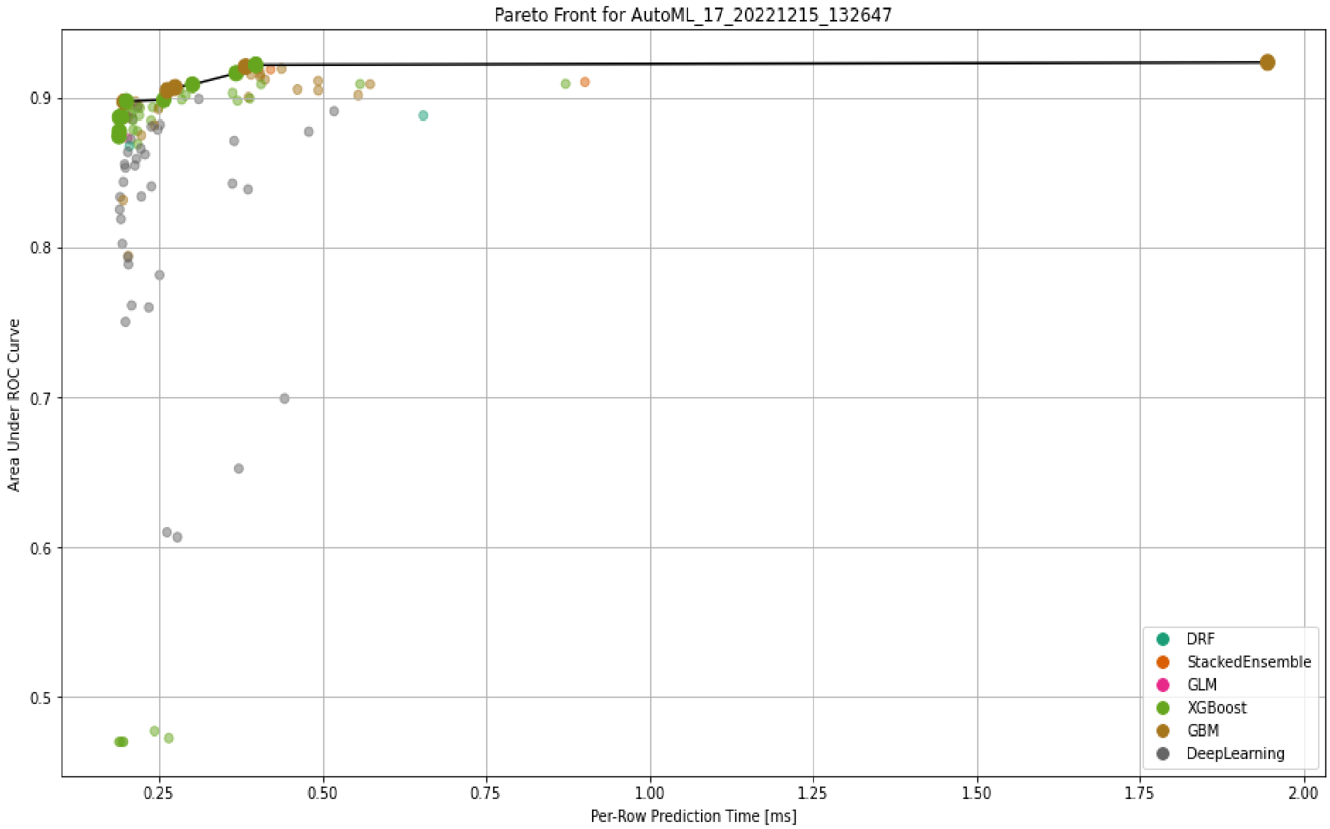


**Figure S3, Pareto Front plot for the 'automl' machine learning.**

### Table S2, metabolomics measurements difference between three treatment groups

| **metabolomics measurements** | | Cholesterol | Linoleic acid | alpha Tocopherol | Alanine total | Asparagine | Aspartic acid | Citric acid | Fructose total | Fumaric acid | Galactose MP | Glucose total | Glutamic acid total |
|---|---|---|---|---|---|---|---|---|---|---|---|---|---|
| **group difference t-test p-value** | T K difference | 0.5866 | *0.0228 | 0.2429 | 0.0508 | 0.0505 | 0.5965 | 0.1510 | *0.0271 | *0.0454 | 0.2456 | 0.1194 | 0.1764 |
| | E K difference | 0.9787 | *0.0482 | **0.0002 | 0.4800 | 0.0930 | 0.7357 | 0.4731 | 0.3565 | 0.6971 | 0.4624 | 0.6782 | 0.4918 |
| | E T difference | 0.5836 | 0.9824 | **0.0045 | 0.2529 | 0.9365 | 0.3720 | 0.5003 | 0.2438 | 0.1550 | 0.6530 | 0.3059 | 0.5151 |
| | | Glutamine | Glutaric acid | Glycine total | Isoleucine total | Lactic acid | Leucine total | Lysine | Malic acid | Methionine total | Ornithine total | Palmitic acid | Phenylalanine total |
| **group difference t-test p-value** | T K difference | 0.1572 | 0.3833 | 0.4904 | 0.7324 | 0.7725 | 0.1793 | 0.1308 | 0.9428 | *0.0385 | 0.1673 | *0.0178 | 0.9470 |
| | E K difference | 0.2885 | 0.7737 | 0.2647 | 0.7144 | 0.5887 | 0.9598 | 0.1941 | 0.3051 | **0.0041 | 0.1322 | *0.0186 | 0.3081 |
| | E T difference | 0.7369 | 0.5770 | 0.5674 | 0.9602 | 0.7665 | 0.2123 | 0.9688 | 0.3003 | 0.2391 | 0.8190 | 0.8413 | 0.2866 |
| | | Proline | Proline total | Pyruvic acid | Serine total | Succinic acid | Threitol | Threonic acid | Threonine total | Tyrosine total | Valine total | myo Inositol | |
| **group difference t-test p-value** | T K difference | 0.0567 | 0.1985 | 0.2136 | 0.8242 | *0.0105 | *0.0224 | 0.9119 | 0.4194 | *0.0456 | 0.5737 | 0.0553 | |
| | E K difference | *0.0164 | 0.0134 | 0.1534 | 0.1608 | *0.0123 | 0.5425 | 0.6703 | 0.1157 | *0.0293 | 0.8457 | 0.5948 | |
| | E T difference | 0.5348 | 0.2129 | 0.7823 | 0.0846 | 0.6286 | 0.1491 | 0.6031 | 0.3905 | 0.6721 | 0.7328 | *0.0138 | |

hsa01100 Metabolic pathways - Homo sapiens (human) (8)

hsa05230 Central carbon metabolism in cancer - Homo sapiens (human) (6)

hsa01230 Biosynthesis of amino acids - Homo sapiens (human) (6)

hsa01210 2-Oxocarboxylic acid metabolism - Homo sapiens (human) (5)

hsa02010 ABC transporters - Homo sapiens (human) (5)

hsa00470 D-Amino acid metabolism - Homo sapiens (human) (4)

hsa01240 Biosynthesis of cofactors - Homo sapiens (human) (4)

hsa01200 Carbon metabolism - Homo sapiens (human) (4)

hsa00330 Arginine and proline metabolism - Homo sapiens (human) (3)

hsa00970 Aminoacyl-tRNA biosynthesis - Homo sapiens (human) (3)

hsa00630 Glyoxylate and dicarboxylate metabolism - Homo sapiens (human) (3)

hsa00770 Pantothenate and CoA biosynthesis - Homo sapiens (human) (3)

hsa00250 Alanine, aspartate and glutamate metabolism - Homo sapiens (human) (3)

**Figure S4, metabolites-pathway mapping results of the eight most valued metabolites in the classifiers results using KEGG mapping tool.**

### Search Grid

**Autoencoder + deeplearning** : activation:[tanh rectifier],
**hidden layers:[(25,25),(12,12),(18,18),(50),(80),(10,10,10),(15,15,15),(20,20,20),(8,8,8,8),(12,12,12,12),(16,16,16,16)]**,
l1:[1e-4,1e-3,1e-2], l2:[1e-4,1e-3,1e-2],
input_drop_out:[0.05,0.1,0.2],hidden_drop_out:[0.3,0.5,0.7],
train_samples_per_iteration:[0,-2],mometum_start:[0,0.5],
rho:[0.5,0.99],quatile_alpha:[0,1]

**XGBoosting:** booster:[gbtree,dart],col_sample_rate:[0.6,0.8,1.0],
col_sample_rate_per_tree:[0.7,0.8,0,9,1.0],max_depth:[5,10,15,20],
min_rows:[0.01,0.1,1,3,5,15,20],reg_alpha:[0.001,0.01,0.1,1,10,100],
reg_lambda:[0.001,0.01,0.1,0.5,1],sample_rate:[0.6,0.8,1]

**GBM:** col_sample_rate:[0.4,0.7,1.0], col_sample_rate_per_tree:[0.4,0.7,1.0],
max_depth:[3--17],min_rows:[1,5,10,15,30,100],
mi_split_improvement:[1e-4,1e-5],sample_rate:[0.5,0.6,0.7,0.8,0.9,1.0]

**Figure S5, machine learning grid search scope.**

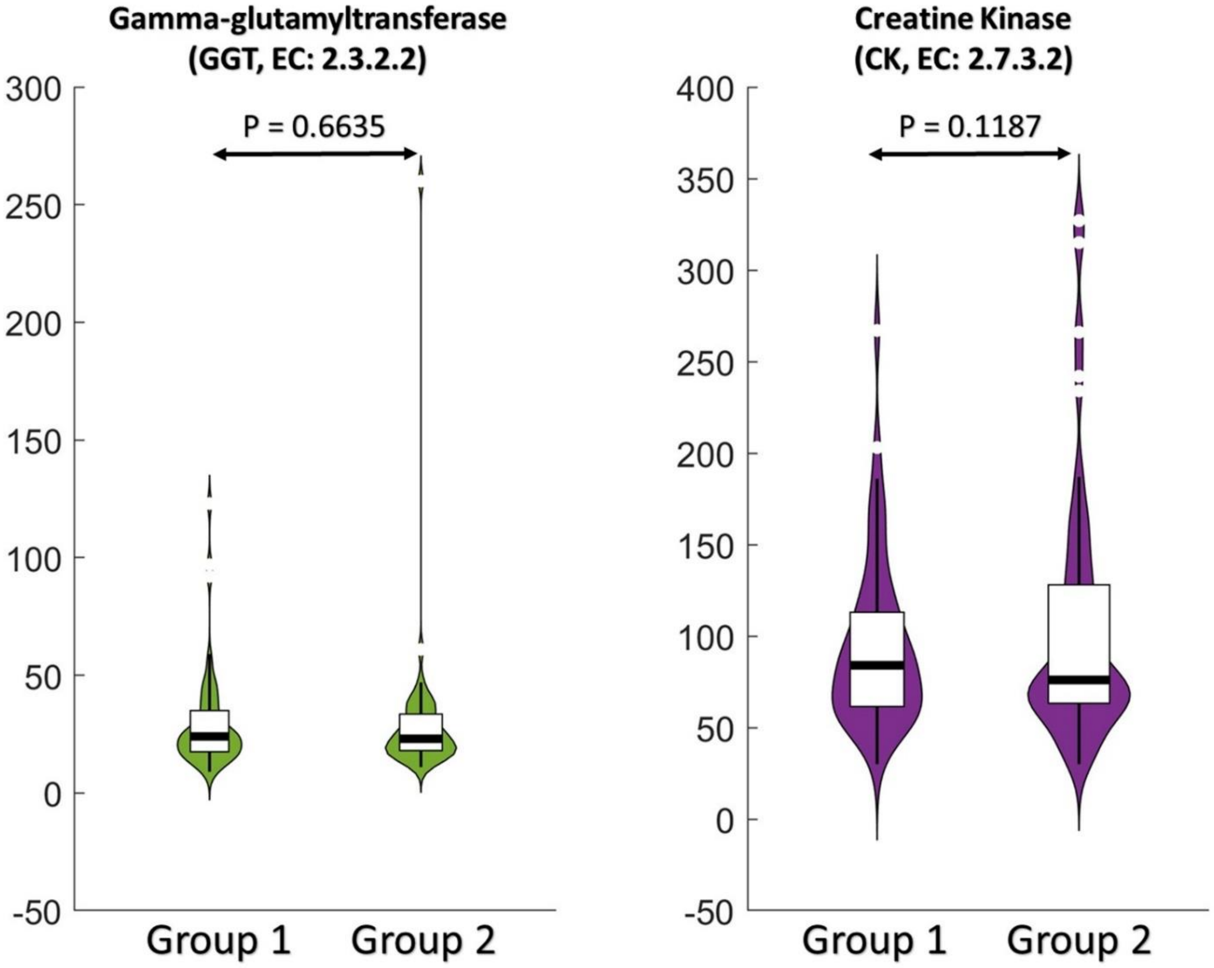


**Figure S6, Comparison of the other two enzymes (left: Gamma-glutamyltransferase, right: Creatine Kinase) between two groups of the first time measurements.**

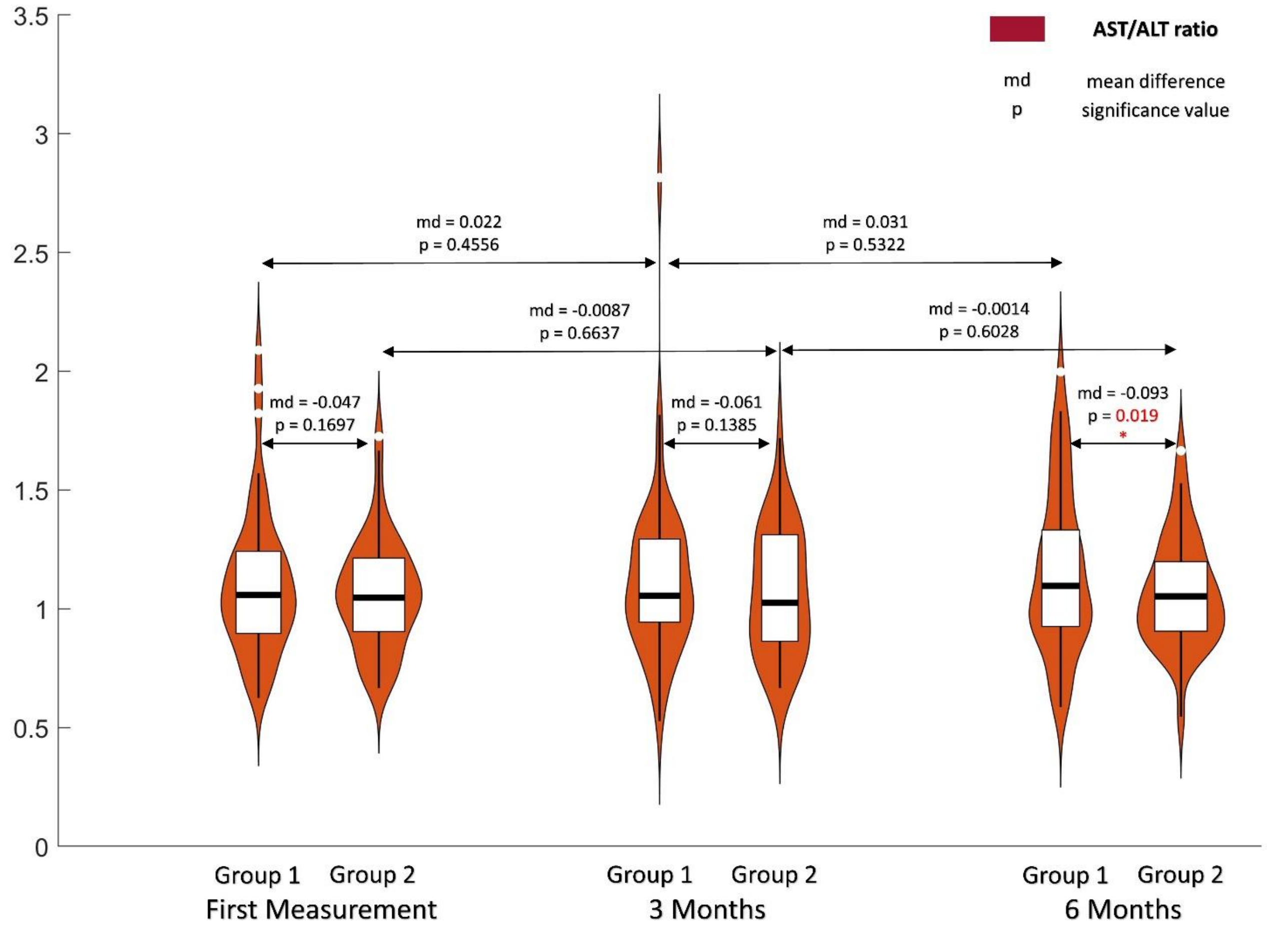


**Figure S7, Comparison of the AST/ALT ratio between two groups over the three time points' measurements.**

### Regression loss based inverse Jacobian analysis

In the inverse Jacobian approach, the Lyapunov Equation (3) is solved using optimization with covariance matrix C and sampled fluctuation matrix D. This linear equation can be rewritten in the form

$$\mathrm{A}_h\, \mathrm{q}_h = \mathrm{b}_h$$

$$\mathrm{A}_d\, \mathrm{q}_d = \mathrm{b}_d.$$

$$(S1)$$

Li, et al. 2023 claimed that under numerical variations in $\mathrm{b}$ the variation of the regression solution $\mathrm{q}$ is much larger compared to the variation in the regression loss $r$. Based on this property, we construct a "regression loss matrix" $\boldsymbol{R}^*$ that aims to capture the relative importance of individual elements in the differential Jacobian, rather than directly calculating the real differential Jacobian matrix. The overall result regression loss matrix $\boldsymbol{R}^*$ is calculated as following

$$q_s^* = \ ({A_c}^T A_c)^{-1} {A_c}^T b_s$$

$$R_{ij}^* = \min_{b_s} \| b_s - A_c q_s^* \|$$

$$(S2)$$

Where $A_c$ is calculated by combining $A_h$ and $A_d$ in Equation (S1) with additional constraint that only that single element $J_{ij}$ may differ between the Jacobians $J_h$ and $J_d$, while all other elements are equal. Because we only have the structure information of the fluctuation matrixes $D_h$ and $D_d$, we generate several (1000 in this article ) samples over possible values for the fluctuation matrixes $b_s = [b_h; b_d]$, following the deduced D structure (related *b* structure in Equation (S1)). Then, $R_{ij}^*$ is calculated as the minimum loss over these samples. In the resulting $\boldsymbol{R}^*$ matrix, larger values indicate correspondingly larger values in the real differential Jacobian matrix. The details of the inverse differential Jacobian algorithm are presented in the original research (Li, et al., 2023).

**Group Jacobian analysis**

Assume that the Jacobian matrix of a N-compounds biological system has p parameters $\alpha = (\alpha_1, \alpha_2, \cdots, \alpha_p)$. The study group contains n objects with Jacobian matrix $Jac\ (\alpha^i)$; for each object, we have $s_i, i \in (1,2 \ldots n)$ samples. The covariance matrix for the samples of each object is $(C_1 \ldots C_n)$. The theoretical fluctuation matrix is D. Then, according to Eq.3,

$$Jac(\alpha^i) * C_i + C_i * Jac(\alpha^i)^T = D_i = D + \Delta D_i, \ \ , i \in (1,2, \ldots, n)$$

Rewrite the equations following,

$$A(\alpha^i) * c_i = d + \Delta d_i, \ \ , i \in (1,2, \ldots, n)$$

In which $c_i$ is a vector of all the independent variables in $C_i$, it has the size of $(\frac{N*(N+1)}{2}, 1)$. Matrix $A(\alpha^i)$ is composed of components in $Jac(\alpha^i)$; it has the size of $(\frac{N*(N+1)}{2}, \frac{N*(N+1)}{2})$. $Vectors\ \ d\ and\ \Delta d_i$ are all the independent variables in $D$ and $\Delta D_i$; it has the size of $(\frac{N*(N+1)}{2}, 1)$. Vector $c_i$ can be calculated as below,

$$c_i = A(\alpha^i)^{-1} * d + \Delta_i \triangleq F(\alpha^i) + \Delta_i$$

In which, A is a continuous function of $\alpha^i$; thus, $F$ is also a continuous function of $\alpha^i$. Now, we regard all the samples as one study group; then, since the parameters are close, we can assume that the steady-state values ${X_0}^i$ are similar. Thus, the covariance matrix of the study group can be estimated as below,

$$c = \frac{\sum_i s_i * c_i}{\sum_i s_i} = \frac{\sum_i s_i * F(\boldsymbol{\alpha^i})}{\sum_i s_i} + \frac{\sum_i s_i * \Delta_i}{\sum_i s_i} = \frac{\sum_i s_i * F(\boldsymbol{\alpha^i})}{\sum_i s_i} + \bar{\Delta}$$

Since $F$ is continuous function of $\alpha^i$, there exist a specific parameter $\alpha^*$ satisfying,

$$c = \frac{\sum_i s_i * F(\boldsymbol{\alpha^i})}{\sum_i s_i} + \bar{\Delta} = F(\boldsymbol{\alpha}^*) + \bar{\Delta}$$

Here the $\alpha^*$ is in the scope of $\alpha^i$. Thus, we derived that one can use a group Jacobian matrix $Jac(\alpha^*)$ to represent all the samples in the group. To test this idea, we utilize carbohydrate energy metabolism (Nazaret and Mazat, 2008) as evaluation model with both evaluation approaches.

Figure S5 demonstrates the inverse differential results with the first evaluation approach. In this evaluation, we carried out a more general test for continuously changing Jacobian-matrix models. Here, we define the group covariance as the mean value of the covariance for samples from each model in one group. We first generate 2*h models with Jacobian matrix step changing from wild type Jacobian matrix J1 to our generated second conditional model J2: $Jac_i = \frac{i-1}{2*h-1} * J1 + \frac{2*h-i}{2*h-1} * J2$. We define the first group consists of the first h models with Jacobian matrix $(Jac_1 ... Jac_h)$, and the other group are the other h models with Jacobian matrix $(Jac_{h+1} ... Jac_{2h})$. For each model, based on the first evaluation approach, we generated the covariance matrix and then calculated the group covariance matrix. With the two group Jacobian matrixes and the Jacobian matrix structure, we applied our new inverse method. With setting h=10, Figure S8 validates the group Jacobian approach. The left subplot is the exact differential Jacobian matrix between $J1 \text{ and } J2$; the other three subplots present three testing attempts through the method above. We can find all the differential Jacobian components when the sample number is enough in the top list, though the regression loss differences between correct components and wrong components are not so clear.

Moreover, we test the group Jacobian approach with not enough number of samples. For each condition 'h' and 'd', we applied random perturbation (max. 10%) to the reaction rate parameter in reaction $R2: Pyrute + NADHc \rightarrow Lactose + NADc$ to generate a sample model in this group. Then we utilize SDE simulation to generate a sample for this model. For each condition, we generate 100 samples through the approach above and calculated the group covariance matrix of these samples. Together with the Jacobian matrix structure, we test our inverse differential Jacobian approach. Figure S8 illustrates the results. The left subplot is the exact differential Jacobian matrix; the other three subplots present three testing attempts for this evaluation. Similar to the results in the previous part, the inverse approach can't reach the perfect differential Jacobian anymore, but still, we can find several differential Jacobian components in the top list.

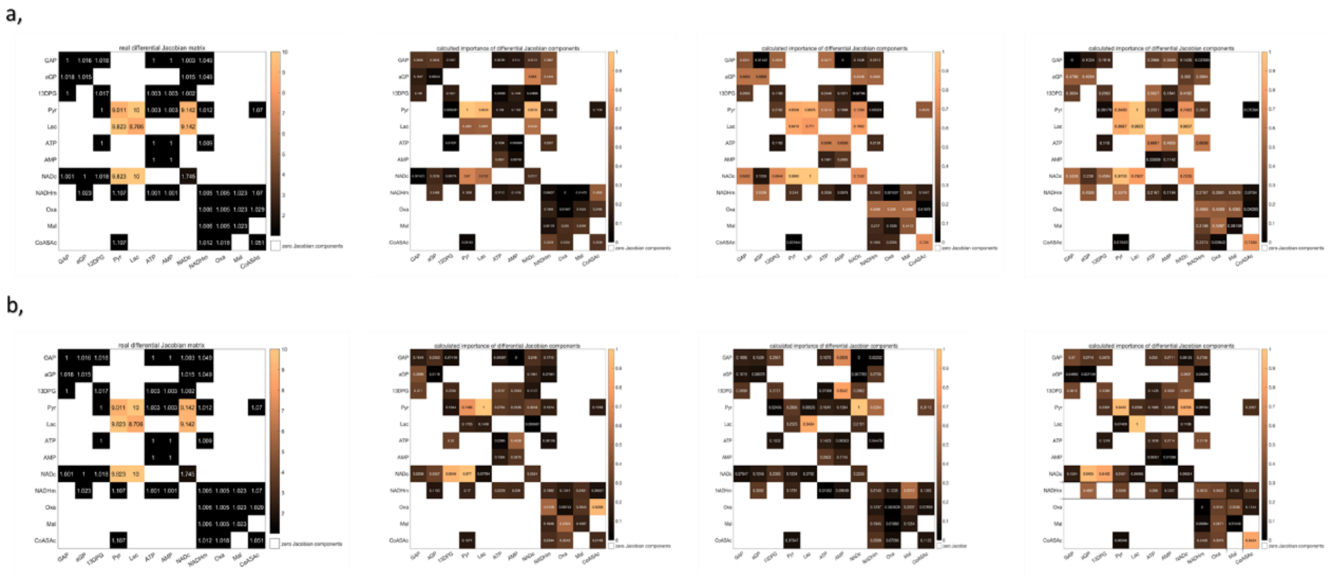

**Figure S8. The inverse differential Jacobian analysis results for the group Jacobian test**. The test contains both evaluation approach: a, the first evaluation approach with enough sample numbers; b, the second evaluation approach with 100 samples from SDE simulation.

### Reference


Li, J., Waldherr, S. and Weckwerth, W. COVRECON: Automated Integration of Genome- and Metabolome- Scale Network Reconstruction and Data-driven Inverse Modeling of Metabolic Interaction Neworks. *Bioinformatics* 2023.

Nazaret, C. and Mazat, J.-P. An old paper revisited:"A mathematical model of carbohydrate energy metabolism. Interaction between glycolysis, the Krebs cycle and the H-transporting shuttles at varying ATPases load" by VV Dynnik, R. Heinrich and EE Sel'kov. *Journal of theoretical biology* 2008;252(3):520-529.

Oesen, S., *et al.* Effects of elastic band resistance training and nutritional supplementation on physical performance of institutionalised elderly—A randomized controlled trial. *Experimental gerontology* 2015;72:99-108.
